# Functional modelling of human infertility-associated genetic variants using *C. elegans* and mice

**DOI:** 10.64898/2026.09.30.755673

**Authors:** Omar El Tokhy, Euan Mitchell, Consuelo Barroso, Jennifer Lawson, Owen R. Davies, Channa N. Jayasena, Joris A. Veltman, Ian R. Adams, Enrique Martinez-Perez

## Abstract

Infertility affects one in six couples worldwide and has a significant genetic component. Genome sequencing efforts have identified increasing numbers of potentially pathogenic variants, including in conserved meiotic genes. However, the lack of functional validation remains a major barrier to establishing variant pathogenicity and clinical diagnoses. Here, we establish an experimental pipeline integrating *C. elegans* and mouse in vivo models with in silico protein structure predictions to functionally validate potentially pathogenic infertility-associated variants. Using CRISPR-Cas9 we generated worm and mouse models of a homozygous missense variant (P638L) in MSH4, a mismatch repair protein essential for meiotic recombination, identified in a patient with non-obstructive azoospermia (NOA). Homozygous *msh-4^P546L^* worms exhibited severe meiotic defects including an absence of chiasmata, accumulation of DNA damage, and loss of crossover markers. Comparable defects were observed in spermatocytes of homozygous *Msh4^P660L/P660L^* mice, while *Msh4^P660L/P660L^* ovaries lacked oocyte-containing follicles, demonstrating that this variant causes infertility in both sexes. Together, these findings support a model in which the MSH4 P638L substitution disrupts the conversion of early recombination events into crossovers, leading to NOA in men and primary ovarian insufficiency in women. Further worm modelling of human *MSH4* genetic variants identified exclusively in heterozygosity (G649R) or also in homozygosity (Y589C) demonstrates the potential of worm models to predict the pathogenicity of missense variants in meiotic proteins found in the human population. We propose that this integrated worm-mouse pipeline provides a powerful framework for accelerating the genetic diagnosis of infertility and informing the clinical management of affected individuals.

## Introduction

Infertility is a major global health problem, affecting approximately 17% of adults during their lifetime ^1^. Both male and female factors contribute substantially to infertility, and genetic defects are increasingly recognised as important causes of impaired reproductive function. In females, primary ovarian insufficiency (POI), a disorder involving premature loss of normal ovarian activity, is a leading cause of total or partial loss of fertility and affects approximately 1 in 100 women under the age of forty ^2^. In males, the most severe form of infertility is non-obstructive azoospermia (NOA), characterised by a complete absence of sperm in the ejaculate. Recent studies reveal a genetic origin for a large proportion of NOA cases ^1,3^. Accordingly, increasing numbers of genetic variants in testis-expressed genes have been described in men with NOA using whole genome and exome sequencing (WES/WGS) ^4–7^. Despite this, few of these variants have undergone functional validation to prove disease causation ^8^. In the absence of efficient systems to screen the pathogenicity of the increasingly large number of NOA-associated variants, such as cell line models for human spermatogenesis, most NOA patients are currently classified as idiopathic and remain untreatable. Thus, there is a need to functionally validate variant-disease relationships both for genetic diagnosis and to gain understanding of the mechanistic basis of male infertility.

NOA is related to defects in spermatogenesis, a complex developmental process that involves mitotic proliferation of diploid spermatogonia, followed by meiosis to reduce the ploidy of diploid spermatocytes to haploid, and finally morphological maturation of post-meiotic haploid spermatids into mature sperm ^9^. Meiosis itself involves two consecutive rounds of chromosome segregation following a single round of DNA replication ^10^. Early meiotic stages are characterised by substantial changes in chromosome structure driven by meiosis-specific cohesin complexes that serve as a scaffold for the assembly of proteinaceous axial elements needed for pairing and recombination between homologous chromosomes (homologues) ^11^. These processes involve the assembly of the synaptonemal complex (SC) between paired homologues, the deliberate formation of DNA double strand breaks (DSBs) to initiate recombination, and the repair of a subset of these DSBs as inter-homologue crossover (CO) events ^12^. COs, together with sister chromatid cohesion provide the basis of chiasmata, physical connections that allow the correct orientation of the homologues on the first meiotic spindle ^10^. Quality control of pairing and recombination is a key aspect of sperm formation in mice, as defects in these processes trigger apoptosis of the defective spermatocytes ^13^. This is best exemplified by the lack of sperm in mouse mutants carrying null mutations on meiotic genes required for pairing and recombination ^14,15^. Notably variants in a number of meiotic genes, including those involved in pairing and recombination, have been identified as causative for NOA, with spermatogenesis in these patients often experiencing meiotic arrest and death of defective meiotic spermatocytes ^16^. As most proteins required for key meiotic prophase events show good evolutionary conservation, findings from model organisms amenable to genetic manipulation of the meiotic machinery, such as mouse and the nematode *C. elegans*, can provide key insights into the mechanistic basis of NOA.

The MutS protein Homolog 4 (MSH4) is an evolutionarily conserved protein required for normal levels of CO formation during meiosis in yeast, *C. elegans* and mouse ^17–19^. Together with MSH5, MSH4 functions as a heterodimer that binds and stabilises double Holliday junction (dHJ) recombination intermediates, thereby enabling the loading of the MLH1-MLH3 complex at these sites to promote the resolution of these intermediates as CO events ^20–22^. *Msh4* is essential for progression through meiosis in mice, with *Msh4*^-/-^ knockout male mice showing severe defects in chromosome pairing and synapsis that causes a zygotene-like arrest during meiotic prophase and completely ablates production of mature spermatozoa ^18^. This arrest is likely caused by defects in the stabilisation of dHJ intermediates, causing synapsis defects that perturb meiotic sex chromosome inactivation and sex body formation, which ultimately leads to pachytene cell death ^13,23^. Oocytes of *Msh4*^-/-^ knockout female mice also exhibit defects in chromosome synapsis and oocyte loss during meiotic prophase ^18^. Similar phenotypes are observed in *Msh5*^-/-^ male and female mutant mice ^24,25^. Interestingly, mice homozygous for mutations in the ATPase domain of MSH5 (*Msh5^GA/GA^*) display metaphase I arrest instead of the earlier meiotic prophase arrest observed in *Msh5*^-/-^ null mutants ^24–26^. Thus, although mutating the ATPase domain of MSH5 does not appear to prevent the ability of MSH5 to promote synapsis, it appears to hinder the ability of MSH5 to ensure chiasma formation and therefore homologue alignment during metaphase I. Thus, MSH4-MSH5 likely plays key roles in both stabilising meiotic recombination intermediates required for synapsis and in promoting CO formation during mouse meiosis.

The essential requirement for MSH4-MSH5 during meiosis likely also extends to humans. Genetic variants in *MSH4* and *MSH5* are associated with NOA in men ^27–31^ and with POI in women ^30–32^. Notably, Krausz *et al*. (2020) identified a novel homozygous recessive missense variant in *MSH4* (P638L) in a NOA patient in which testis biopsies showed that spermatocytes progressed to metaphase I before undergoing cell death. This is a significantly later meiotic arrest than that reported for *Msh4*^-/-^ null mice ^18^, but is similar to the metaphase I arrest observe in homozygous *Msh5^GA/GA^* (ATPase) mouse mutants ^26^. It is not clear whether the metaphase I arrest of the NOA patient homozygous for *MSH4^P638L^* reflects MSH4 P638L having similar effects on MSH4-MSH5 function as the MSH5 GA ATPase mutation, different responses of mouse and human spermatocytes to loss of MSH4 function, or additional genetic variants present in this NOA patient.

Here, we have used CRISPR/Cas9 genome editing to create *C. elegans* and mouse models of the infertility-associated *MSH4^P638L^* variant to determine its impact on key meiotic events. Worm and mouse models of the *MSH4^P638L^* variant display a failure in converting early recombination events into COs, a phenotype reminiscent of *Msh4* null mutants in both systems. Further worm modelling of common and rare *MSH4* genetic variants identified in the human population suggests that the rare variant (G649R) is pathogenic, whereas the common variant (Y589C) is not. The workflow described here represents a pipeline to assess the phenotypic consequences of infertility-associated genetic variants in conserved meiotic genes using worms and mice as model organisms, thus providing a functional assay to determine whether specific genetic variants cause human infertility.

## Results

### The region containing the NOA-associated variant MSH4^P638L^ is conserved in *C. elegans* and mouse

Exome sequencing data from a NOA patient with spermatogenic maturation arrest identified a novel homozygous missense variant in *MSH4* (Pro638Leu, P638L) ^27^. The male patient homozygous for the *MSH4^P638L^* variant shows meiotic metaphase I arrest from immunostaining of testicular biopsies. However, the meiotic arrest phenotype in this patient is later than that reported for *Msh4^-/-^* null mice and the effect of the P638L substitution on MSH4 function is not known. P638 is located between domains IV and V of MSH4 (Figure 1a), it is conserved in mouse MSH4 (P660) and *C. elegans* MSH-4 (P546) (Figure 1b), and *AlphaFold 3* modelling places it in a structured interdomain region of the human MSH4-MSH5 heterodimer (Figure 1c and Figure S1). In this model, the backbone dihedral angles of P638 fall within the generally favoured region of the Ramachandran plot, indicating that the local backbone conformation is not proline-specific (Figure 1d). However, the proline side chain is tightly packed within a hydrophobic pocket formed by surrounding MSH4 residues (Figure 1e), which is unlikely to accommodate the longer leucine side chain. Hence, P638L would likely create steric clashes within this pocket, so is predicted to interfere with local MSH4 folding or stability and/or induce a conformational change that compromises MSH4 function. On this basis, and given the evolutionary conservation of MSH4 P638, we decided to investigate the functional relevance of the P638L substitution identified in this infertile male patient by creating *C. elegans* and mouse models. To do so, we first used CRISPR/Cas9 genome editing to introduce the corresponding substitution (P546L) into *him-14*, the *C. elegans* orthologue of human *MSH4* (hereafter referred to as *msh-4*).

**Figure 1.**
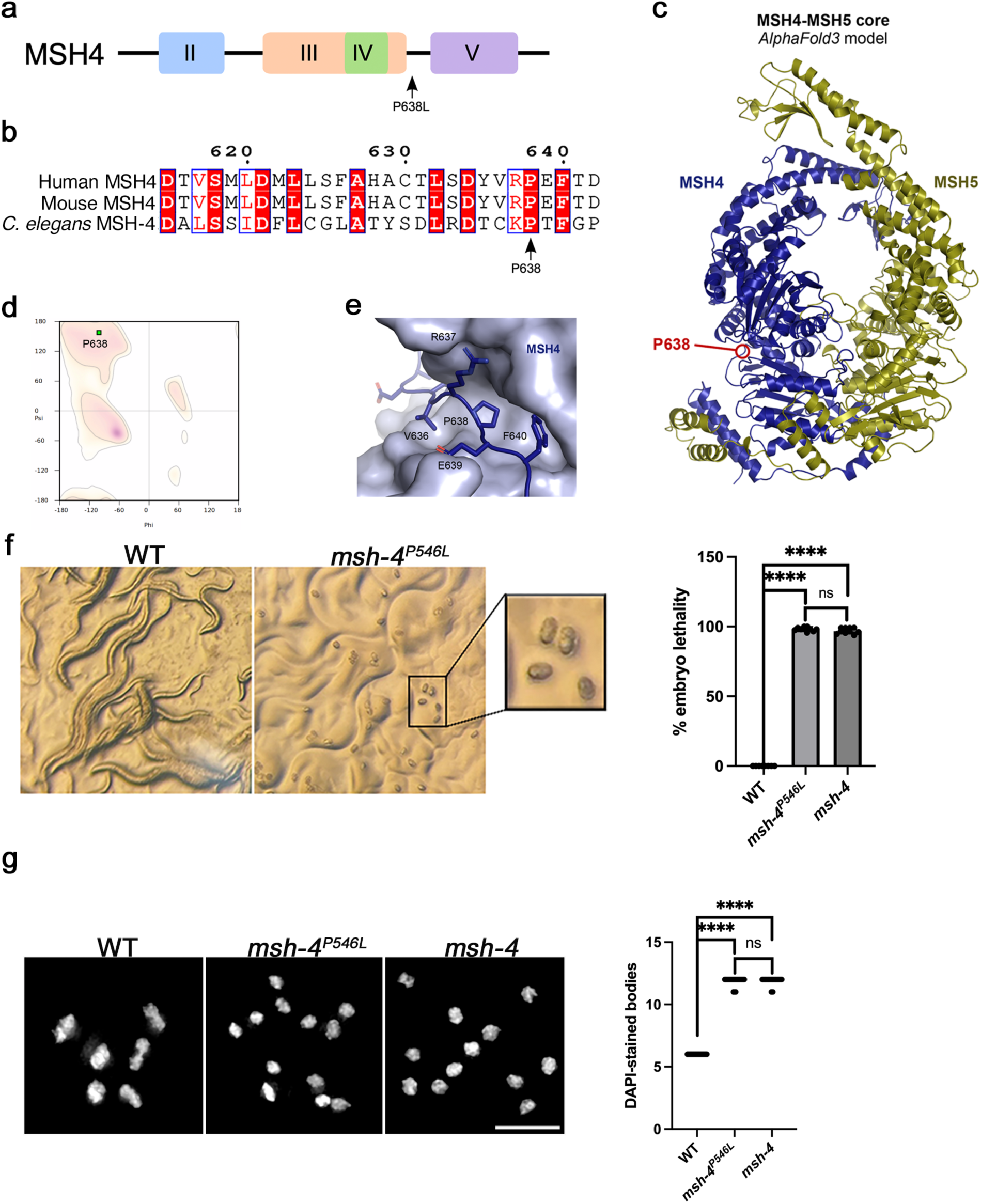
*C. elegans msh-4^P546L^* mutants display fertility defects. **(a)** Diagram of the MSH4 protein indicating domains II-V and the location of the P638L NOA variant. **(b)** Alignment of the MSH4 region containing P638 in human, mouse, and *C. elegans*. Numbering refers to the human protein. **(c)** *AlphaFold 3* model of the structure core of a human MSH4-MSH5 1:1 complex (MSH4, amino acids 148-926, blue; MSH5, amino acids 47-834, yellow) showing the location of amino acid P638. **(d)** Ramachandran plot showing that the predicted dihedral angles of amino acid P638 are within the favoured region for general amino acids. **(e)** Structural environment of P638 within the model, showing how local side-chains are predicted to pack against the remainder of the MSH4 surface. **(f)** Images from plates containing progeny from a single wild type control hermaphrodite or a single *msh-4^P546L^* mutant hermaphrodite 2 days after removal of the mother. Note the presence of larvae in the wild type plate, while the *msh-4^P546L^* mutant plate only contains unhatched (dead) embryos. Graph shows quantification of the percentage of dead embryos amongst the progeny of worms of indicated genotypes (progeny from between 8 and 10 worms scored per genotype, error bars indicate mean with 95% CI, p values were calculated using a two-tailed Mann-Whitney U test (**** indicates p < 0.0001). **(g)** Projections of nuclei from diakinesis oocytes stained with DAPI, note the presence of 6 DAPI-stained bodies in the wild type nucleus and of 12 DAPI-stained bodies in the *msh-4^P546L^* mutant nucleus. Graph shows quantification of the number of DAPI-stained bodies in diakinesis oocytes of indicated genotypes (30 nuclei scored per genotype), error bars indicate mean with 95% CI, p values were calculated using a two-tailed Mann-Whitney U test (**** indicates p < 0.0001). Scale bar = 5 µm.

### *C. elegans msh-4^P546L^* mutants modelling the *MSH4^P638L^* human mutation display severe meiotic and fertility defects

The hermaphrodite *C. elegans* germline produces a fixed complement of sperm during early larval development, which is stored in the spermatheca, before permanently switching to oogenesis in adult worms. Fertilisation takes place as oocytes pass through the spermatheca after completing meiotic prophase. Worm mutants with defects in meiotic recombination frequently mis-segregate their chromosomes during meiosis and therefore self-fertilisation creates mostly aneuploid embryos unable to complete normal development and hatch into viable larvae. Therefore, the presence of a high percentage of unhatched dead embryos is an expected phenotype of recombination deficient worm mutants ^33^. As embryonic development in *C. elegans* takes approximately 14 hours, we scored the frequency of unhatched embryos amongst the progeny of *msh-4^P546L^* mutants 24 hours after the embryos were laid. In plates containing wild-type (WT) mothers, nearly 100% of embryos hatch, whereas most embryos (98%) fail to hatch in *msh-4^P546L^* mutants, as also observed in *msh-4* null mutants used as a control (Figure 1f). To determine if the high level of embryonic lethality of *msh-4^P546L^* mutants is due to defects in meiotic recombination, we assessed chiasma formation in oocytes at the diakinesis stage, when chromosome condensation and the large volume of the oocyte nucleus facilitate visualisation of individual chromosomes. The *C. elegans* genome comprises six pairs of homologous chromosomes, therefore diakinesis oocytes of wild type worms contain six DAPI-stained bodies, with each one corresponding to a pair of homologous chromosomes attached by a chiasma (bivalent) (Figure 1g). In contrast, diakinesis oocytes of recombination mutants display up to 12 DAPI-stained bodies, with each one corresponding to a single chromosome (univalent). Most diakinesis oocytes of *msh-4^P546L^* and *msh-4* null mutants contained twelve DAPI-stained-bodies, demonstrating a near complete absence of chiasmata (Figure 1g). These results demonstrate the presence of a severe recombination defect in *msh-4^P546L^* homozygous mutants.

### *C. elegans msh-4^P546L^* homozygous mutants fail to convert early recombination intermediates into crossovers

To understand the origin of the chiasma defect of *msh-4^P546L^* mutants, we performed immunostaining experiments to investigate the formation and resolution of early recombination foci containing the DNA repair protein RAD-51, which localises to ssDNA following DSB formation ^34,35^, and the formation of crossover-fated events containing COSA-1 during late pachytene ^36^. We monitored the appearance and disappearance of RAD-51 by dividing the germline into five equal-size regions between the onset of meiotic prophase (leptotene) and the end of late pachytene and quantifying the number of RAD-51 foci in each nucleus of these five zones. In wild type germlines RAD-51 foci peak in zone 3 before dropping in zone 4 and becoming barely detectable in zone 5 (Figure 2a). *msh-4^P546L^* mutant germlines accumulate much higher numbers of RAD-51 foci than wild type controls, especially in zones 3 and 4, a pattern similar to that observed in *msh-4* null mutants ^37^, evidencing an accumulation of early recombination intermediates (Figure 2a). In contrast, late recombination events marked by COSA-1 were largely lacking in *msh-4^P546L^* mutant germlines, in which most nuclei contained no COSA-1 foci. This contrasts with the 6 foci observed in wild type controls (Figure 2b) and is similar to the lack of COSA-1 foci observed in *msh-5* mutants ^36^. This severe reduction in the number of late pachytene COSA-1 foci despite earlier accumulation of RAD-51 foci suggests that the primary defect of *msh-4^P546L^* worm mutants is the inability to convert early recombination intermediates into COs. Nonetheless, as defects in synaptonemal complex assembly also result in accumulation of RAD-51 foci and impaired chiasma formation in *C. elegans* ^37^, we investigated whether *msh-4^P546L^* mutants display synapsis defects. Staining with antibodies against axial element HIM-3 ^38^ and central region component SYP-1 ^39^ demonstrated no obvious synapsis defects in pachytene nuclei of *msh-4^P546L^* mutants (Figure S2), similar to the situation in *msh-4* null mutants ^37^. Therefore, the MSH-4 P546L substitution specifically impairs the conversion of RAD-51 intermediates into late recombination events capable of yielding inter-homologue crossovers, phenocopying defects of *C. elegans msh-4* null mutants.

**Figure 2.**
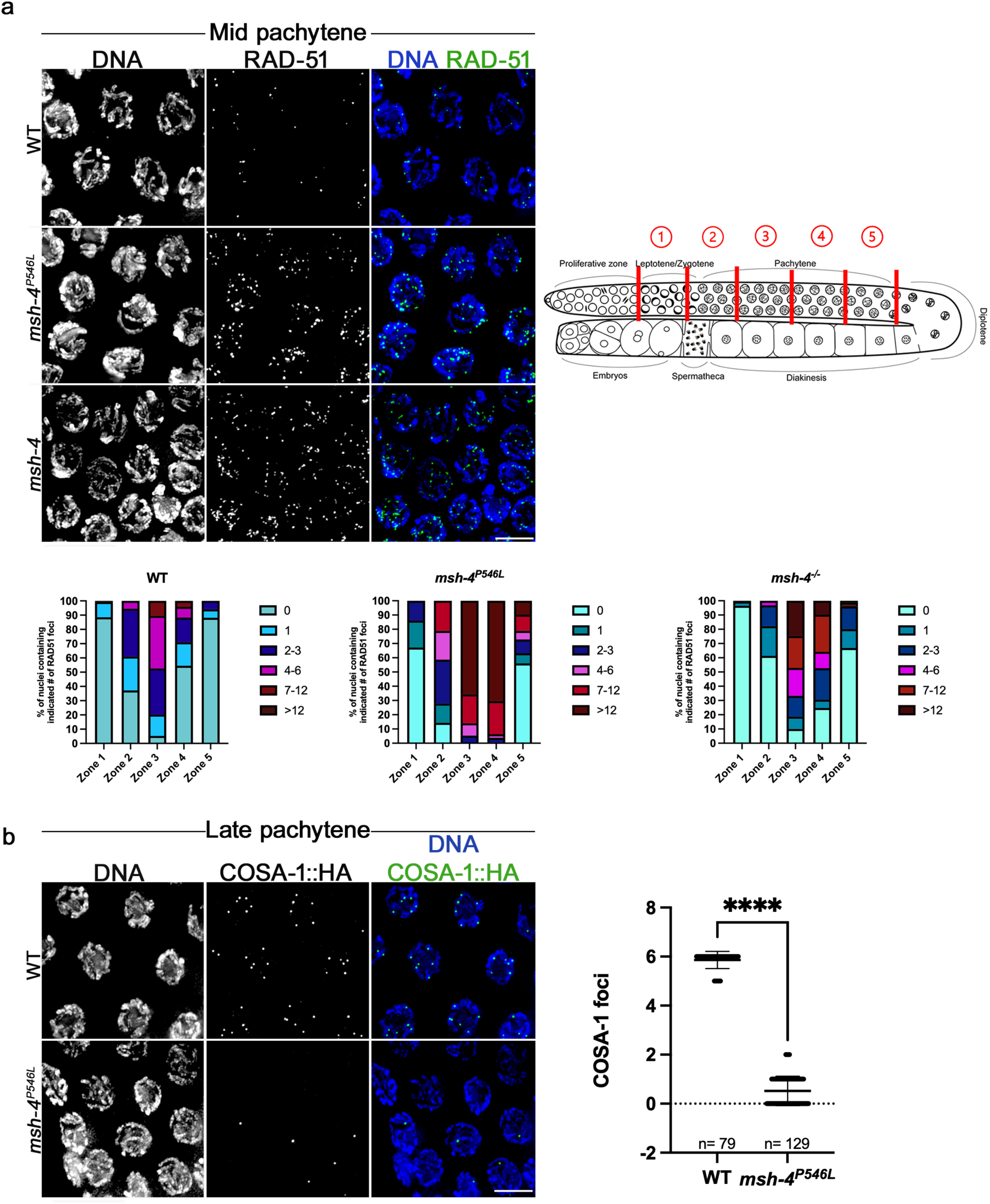
*C. elegans msh-4^P546L^* mutants display meiotic recombination defects. **(a)** Quantification of RAD-51 foci in germlines of indicated genotypes. Images show projections of mid pachytene nuclei stained with anti-RAD-51 antibodies and DAPI. Note that *msh-4^P546L^* and *msh-4* mutants display higher numbers of RAD-51 foci than wild type controls. Cartoon of a dissected *C. elegans* germline indicating how the region between the onset of leptotene and the end of pachytene was divided into five equal-size regions. Graphs show the percentage of nuclei displaying a given number of RAD-51 foci per zone. Three germlines were scored per genotype, total number of nuclei analysed per zone: 130, 163, 132, 121, 118 (WT); 180, 209, 221, 206, 166 (*msh-4^P546L^*); 202, 259, 205, 226, 151 (*msh-4*). **(b)** Projections of late pachytene nuclei of indicated genotype stained with DAPI and anti-HA antibodies (to visualise COSA-1::HA protein), graph shows quantification of COSA-1::HA foci in late pachytene nuclei form three germlines per genotype (n indicates the total number of nuclei analysed per genotype), error bars indicate mean with standard deviation, p value was calculated using a two-tailed Mann-Whitney U test (**** indicates p < 0.0001). Scale bar = 5 µm in all panels.

### *Msh4^P660L/P660L^* mutant mice display strong azoospermic phenotypes

We next investigated whether the MSH4 P638L substitution similarly impairs meiotic recombination and gametogenesis using a mammalian model. To model the *MSH4^P638L^* variant in mice, we used CRISPR/Cas9 gene editing to introduce the equivalent mutation in mouse *Msh4* (P660L) by homology-directed repair in mouse zygotes (Figure S3a). As part of this gene editing strategy, we introduced a synonymous blocking mutation in the CRISPR/Cas9 PAM site to prevent successful targeting events from being re-targeted. In parallel, we also generated control CRISPR/Cas9 gene edited mice with the synonymous PAM site substitution in the absence of the P660L mutation (Figure S3b). These PAM control mice ought to control for any unexpected effects of the PAM site substitution and for off-target effects of CRISPR/Cas9 gene editing. Founder mice carrying *Msh4^P660L^* or *Msh4^PAM^* alleles identified by PCR and sequencing were backcrossed to C57BL/6J mice and the resulting heterozygous mice crossed to generate *Msh4^P660L/P660L^* or *Msh4^PAM/PAM^* homozygotes. Restriction digests of PCR amplified samples from both the *Msh4^P660L^* and *Msh4^PAM^* mouse lines showed the expected band sizes for wild type, heterozygous and homozygous mice (Figure S3c), and sequencing data from *Msh4^P660L/P660L^* and *Msh4^PAM/PAM^* homozygotes confirmed that these alleles contain the desired nucleotide sequence for each genotype with no additional mutations (Figure S3d).

Phenotypic analysis of *Msh4^P660L/P660L^* mice showed that *Msh4^P660L/P660L^* testes are smaller than wild-type controls and have an average 78.6% reduction in weight relative to controls (Figure 3a-b). Furthermore, we could not detect any sperm in the epididymis of *Msh4^P660L/P660L^* males (Figure 3b). Taken together, the gross testis and epididymal sperm count data suggest that the *Msh4* P660L mutation severely disrupts spermatogenesis. In contrast, we could not detect a difference in testis size, weight or epididymal sperm count in *Msh4^PAM/PAM^* homozygotes with the silent PAM substitution relative to wild-type controls (Figure S4a-b). Thus, the disruption of spermatogenesis present in *Msh4^P660L/P660L^* males is likely caused by the P660L amino acid substitution in MSH4.

**Figure 3:**
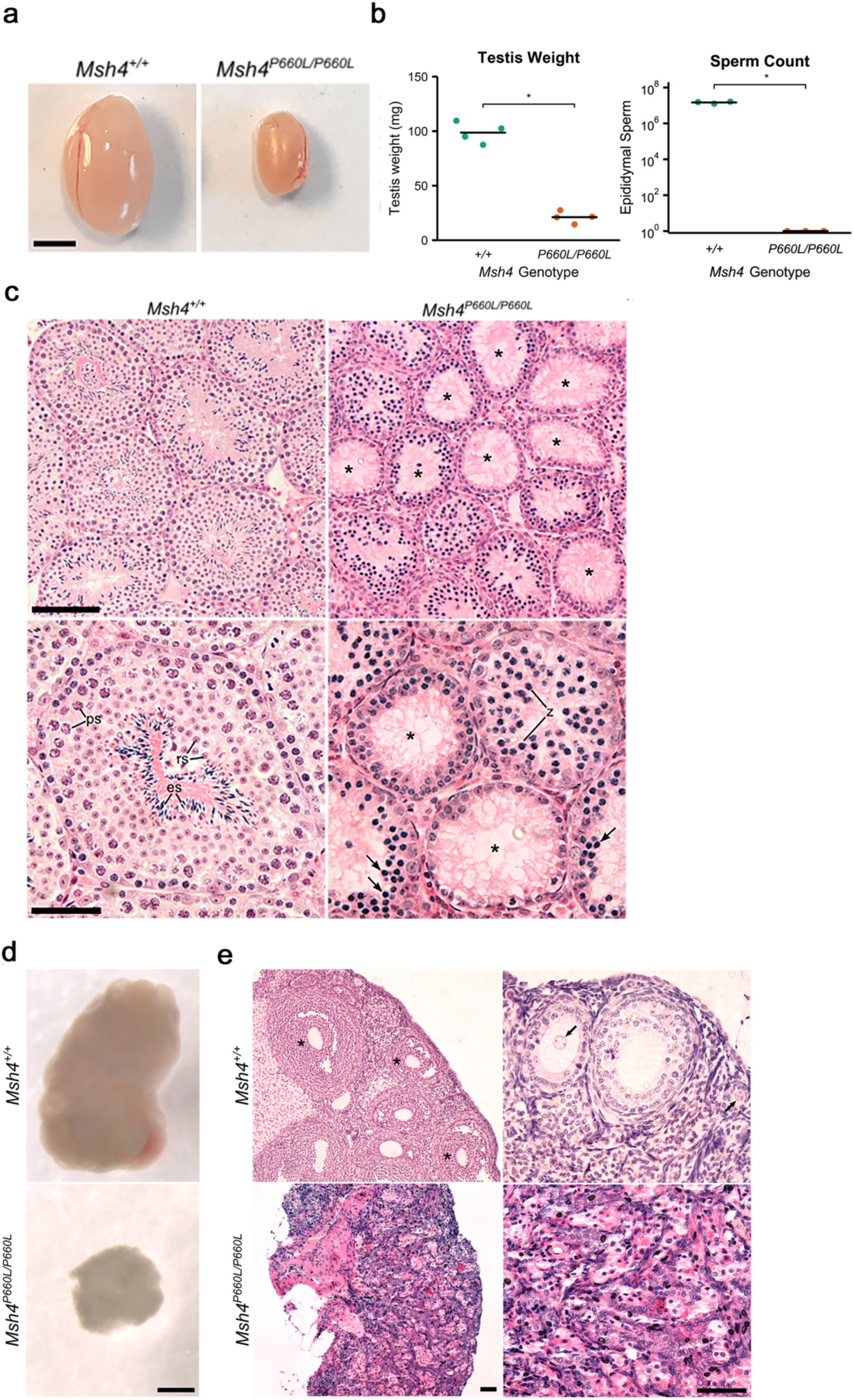
Spermatogenesis and oogenesis are perturbed in *Msh4^P660L/P660L^* mice. **(a)** Example images of *Msh4^+/+^* and *Msh4^P660L/P660L^* testes. Scale bar, 5 mm. **(b)** Testes weights and epididymal sperm counts from *Msh4^+/+^* and *Msh4^P660L/P660L^* mice. Horizontal bars represent means. * = p <0.05 (Student’s t-test). n = 4 mice (testis weight), n=3 mice (sperm count). **(c)** Haematoxylin and eosin staining of testes sections. Asterisks indicate tubules with depletion of post-meiotic spermatids. Examples of main cell types are annotated; ps = pachytene spermatocyte, rs = round spermatid, es = elongated spermatid, z = zygotene-like spermatocyte, arrow = pyknotic cell. Scale bars, top panel= 100 μm and bottom panel= 50 μm. **(d)** Images of *Msh4^+/+^* and *Msh4^P660L/P660L^* ovaries. Scale bar= 500 µm. **(e)** Haematoxylin and eosin staining of *Msh4* ovary sections, showing both overall tissue structure (left) and individual follicles (right). Follicles are indicated with an asterisk in the low magnification image, examples of oocyte nuclei within primary or secondary follicles are indicated with an arrow in the high magnification image. Images are representative of ovaries from 4 mice per genotype. Scale bars= 50 µm.

To investigate the spermatogenesis defect in *Msh4^P660L/P660L^* males further, we analysed testis sections histologically. As expected, all stages of spermatogenesis were detectable in seminiferous tubules within *Msh4^+/+^* testes; however, seminiferous tubules within *Msh4^P660L/P660L^* testes lacked post-meiotic round spermatids and elongated spermatids, and often contained accumulations of dying cells with pyknotic nuclei (Figure 3c). Though meiotic spermatocytes with a zygotene-like nuclear morphology were readily detectable in *Msh4^P660L/P660L^* seminiferous tubules, we could not detect later stages of meiosis (Figure 3c). These defects were not apparent in testis sections from *Msh4^PAM/PAM^* homozygous mice with the silent PAM substitution, which appeared to be grossly normal (Figure S4c). Thus, the *Msh4* P660L mutation appears to cause defects in spermatogenesis and progression through meiotic prophase I in mice. Notably, the lack of any spermatocyte nuclei progressing to pachytene or later indicates that *Msh4^P660L/P660L^* spermatocytes undergo a complete meiotic arrest during prophase I consistent with the *Msh4^-/-^* null mouse phenotype ^18^, rather than the later metaphase I arrest phenotypes reported in the human *MSH4^P638L^* infertility patient ^27^ and *Msh5^GA/GA^* ATPase domain mouse mutant ^26^.

### *Msh4^P660L/P660L^* ovaries have defects in oogenesis and folliculogenesis

Although the *MSH4* P638L variant was originally identified as a cause of male infertility, the *MSH4* P638L index patient was reported to have 2 sisters with infertility ^27^, which could potentially reflect this *MSH4* variant perturbing both oogenesis and spermatogenesis. Given this possibility and other studies identifying *MSH4* variants in human POI patients ^30,32^, we assessed whether the *Msh4^P660L^* variant can cause defects in female meiosis and oogenesis. Ovaries from *Msh4^P660L/P660L^* female mice were visibly smaller than wild-type control ovaries (Figure 3d). Furthermore, we could not detect follicles at any stage of folliculogenesis in *Msh4^P660L/P660L^* ovary histological sections, rather the *Msh4^P660L/P660L^* ovaries appeared to only contain stromal tissue (Figure 3e). No such defects were detected in *Msh4^PAM/PAM^* homozygous ovaries (Figure S4d-e). These defects in oogenesis and folliculogenesis in *Msh4^P660L/P660L^* female mice are similar to those reported in *Msh4^-/-^* null female mice ^18^.

### *Msh4^P660L/P660L^* mice spermatocytes exhibit defects in meiotic chromosome pairing and progression

To investigate the meiotic defects in *Msh4^P660L/P660L^* spermatocytes in more detail, we prepared chromosome spreads from *Msh4^P660L/P660L^* testes and immunostained them with antibodies against the synaptonemal complex components SYCP3 and SYCP1 ^15,40^ to monitor progression through meiotic prophase I.

In the *Msh4^+/+^* chromosome spreads, we observed spermatocytes at leptotene, zygotene, pachytene and diplotene stages of prophase I, with the majority in either pachytene (∼65%) or diplotene (∼26%) (Figure 4a). However, in the *Msh4^P660L/P660L^* homozygotes we were unable to detect any nuclei progressing to or beyond pachytene, and the majority of SYCP3 positive nuclei instead displayed an aberrant zygotene-like phenotype with fully formed SYCP3 axial elements but only short synapsed stretches of SYCP1 that usually did not extend along an entire chromosome axis (Figure 4a). Some SYCP1 staining in these zygotene-like spermatocytes was present as axis-associated foci on both the synapsed and asynapsed chromosome axes (Figure 4a). Furthermore, staining for sex body formation and DNA damage using antibodies to γH2AX ^41,42^ showed reliable formation of sex bodies in the control spermatocytes whereas *Msh4^P660L/P660L^* spermatocytes had γH2AX accumulating along asynapsed chromosome axes and no discernible sex bodies (Figure 4b). The *Msh4^P660L/P660L^* spermatogenesis phenotype is consistent with asynapsed chromosomes causing pachytene cell death through sequestering the transcriptional silencing machinery away from the sex chromosomes and disrupting meiotic sex chromosome inactivation ^13,23^. In contrast, we could not detect any defects in chromosome synapsis or progression through meiosis in chromosome spreads from testes of *Msh4^PAM/PAM^* homozygous mice with the silent PAM nucleotide substitution (Figure S5a). Interestingly, although *Msh4^P660L/P660L^* homozygous mouse spermatocytes arrest meiosis at an earlier stage than MSH4 P638L patient spermatocytes, the failure in sex body formation appears to occur in both species ^27^.

**Figure 4:**
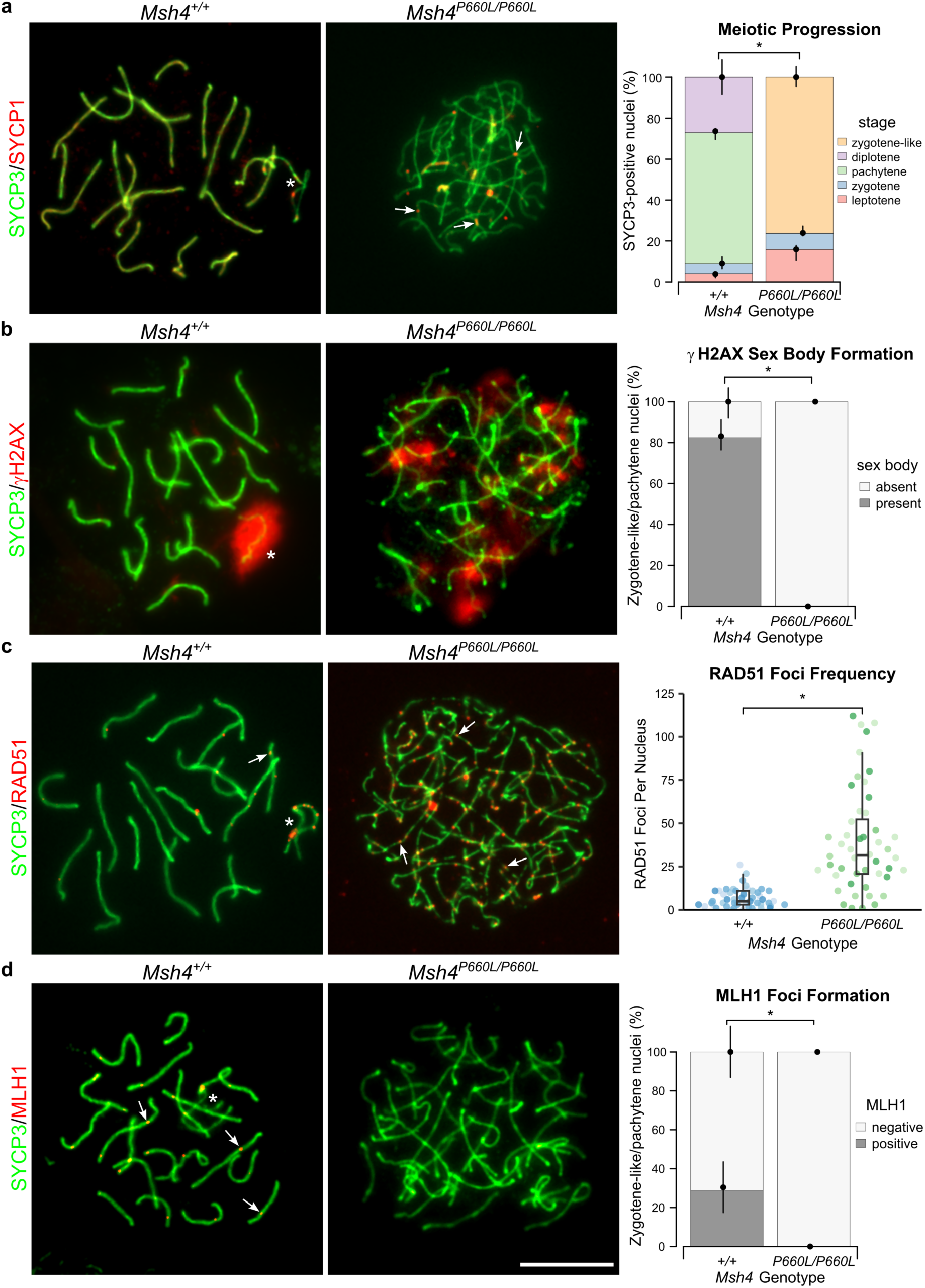
*Msh4^P660L/P660L^* mice show synapsis and recombination defects during meiotic prophase. **(a)** Widefield imaging of pachytene *Msh4^+/+^* and zygotene-like *Msh4^P660L/P660L^* spermatocyte meiotic chromosome spreads immunostained for the synaptonemal complex components SYCP3 (green) and SYCP1 (red), with quantitation of percentage of total SYCP3-positive nuclei observed at each stage of prophase or in a zygotene-like state. Axis-associated SYCP1 foci and short stretches of SYCP1 are indicated with arrows, sex chromosomes with an asterisk. n = 4 mice per genotype, 122 and 139 nuclei scored in total, * PP>0.999 (Bayesian GLMM); circles and vertical lines represent the means and interquartile distances of animals within each genotype. **(b)** Widefield imaging of pachytene *Msh4^+/+^* and zygotene-like *Msh4^P660L/P660L^* meiotic chromosome spreads immunostained for the synaptonemal complex component SYCP3 (red) and the sex body/DNA damage marker γH2AX (green). Quantitation of total SYCP3 positive nuclei with/without sex bodies is shown on the right. Sex bodies indicated with an asterisk. n = 4 mice per genotype, 131 and 126 nuclei scored in total, * PP>0.999 (Bayesian GLMM); circles and vertical lines represent the means and interquartile distances of animals within each genotype. **(c)** Widefield imaging of pachytene *Msh4^+/+^* and zygotene-like *Msh4^P660L/P660L^* spermatocyte meiotic chromosome spreads immunostained for SYCP3 (green) and RAD51 (red), with quantitation (beeswarm plot overlaid with box-and-whiskers) of number of RAD51 foci on chromosome axes. Beeswarm data points from individual mice are shaded differently for each genotype. Examples of RAD51 foci indicated with arrowheads. n = 4 mice per genotype, 48 nuclei scored in total for each genotype, * PP>0.999 (Bayesian GLMM). **(d)** Widefield imaging of pachytene *Msh4^+/+^* and zygotene-like *Msh4^P660L/P660L^* meiotic chromosome spreads immunostained for SYCP3 (red) and MLH1 (green), with quantitation of MLH1 positive (≥ 15 foci) and negative (0 foci) nuclei per genotype. Examples of MLH1 foci indicated with arrowheads. n = 4 mice per genotype, 90 and 142 nuclei scored in total, * PP>0.999 (Bayesian GLMM); circles and vertical lines represent the means and interquartile distances of animals within each genotype. Scale bar, 10 µm.

### *Msh4^P660L/P660L^* mice spermatocytes show defects in progression of meiotic recombination

MSH4 plays an essential role in meiotic recombination and is thought to stabilise inter-homolog joint molecule recombination intermediates to allow them to progress to late recombination foci ^21,43^. We therefore tested whether meiotic recombination was initiating and progressing normally in *Msh4^P660L/P660L^* spermatocytes. Early recombination foci were examined using the marker protein RAD51, a conserved mammalian homolog of RecA recombinase that binds to early leptotene and zygotene meiotic DSB sites and then becomes gradually removed from autosomes as recombination progresses ^35,44^. Consistent with previous studies ^44^, relatively few RAD51 foci were present in the control *Msh4^+/+^* pachytene spermatocytes, however a much higher frequency of RAD51 foci was observed in the *Msh4^P660L/P660L^* zygotene-like nuclei (Figure 4c). This abundance is likely reflective of altered repair of the early recombination foci, possibly due to either lack of chromosome synapsis and/or additional DSB formation along asynapsed axis regions ^18,45^. This high frequency of RAD51 foci in *Msh4^P660L/P660L^* zygotene-like nuclei is consistent with observations in *Msh4^-/-^* null spermatocytes ^18^, and resonates with the high frequency of RAD-51 foci in *C. elegans msh-4^P546L^* and *msh-4^-/-^* null mutants during pachytene.

The DNA mismatch repair protein MLH1 is a well-characterised marker for late-stage meiotic recombination foci that is recruited to chromosome axes in late pachytene, and plays a role in resolution of recombination intermediates as crossovers ^44,46–48^. As expected, approximately 25% of the pachytene nuclei in *Msh4^+/+^* chromosome spreads were MLH1-positive, whereas MLH1-positive nuclei were absent from *Msh4^P660L/P660L^* chromosome spreads (Figure 4d). Thus, meiotic recombination is not progressing to late stages in *Msh4^P660l/P660L^* spermatocytes, likely due to asynapsed chromosome axes triggering pachytene cell death in *Msh4^P660L/P660L^* spermatocytes before they reach the stage that MLH1 can be detected ^13,23^.

Consistent with our observations from the *C. elegans* model, this mouse model suggests that the NOA-associated *MSH4* P638L variant impairs progression of meiotic recombination and generates a similar phenotype to a *Msh4* null allele. Taken together, data from both the *C. elegans* and mouse models suggest that the MSH4 P638L substitution likely severely impairs MSH4 function in humans and is causative for NOA.

### Using *C. elegans* to determine the pathogenicity of *MSH4* variants present in the human population

The results shown above from our worm and mouse models of the *MSH4* P638L variant demonstrate that worms can be used to determine the pathogenicity of infertility-associated variants for which patient data is available. We next investigated the predictive potential of our pipeline to determine the pathogenicity of variants in *MSH4* present in the human population, but for which no patient data is available. To this end, we first selected *MSH4* variants described in the Genome Aggregation Database (gnomAD), the largest and most widely used publicly available collection of population variation from human sequencing data ^49^. We reasoned that variants detected at low frequency in the population and that have only been detected in heterozygosity are more likely to hinder fertility than variants detected at high frequency and that have been reported in homozygosity. We also used AlphaMissense, a deep learning model that uses AlphaFold structural predictions to calculate the pathogenicity score of single amino acid substitutions ^50^, to predict the pathogenicity score (from 0 to 1) of the MSH4 variants selected from gnomAD. Two variants, *MSH4^G649R^* (low frequency (4.96 x 10^-6^), not detected in homozygosity, high pathogenicity score) and *MSH4^Y589C^* (high frequency (9.8 x10^-3^), detected in homozygosity 123 times, low pathogenicity score) were selected for modelling in *C. elegans*.

The *MSH4^G649R^* variant is not known to cause infertility, but is predicted as pathogenic by AlphaMissense with a score of 0.932 and has a low allele frequency. Notably, MSH4^G649R^ lies downstream of P638, within the same predicted structured interdomain region between domains IV and V of MSH4 (Figure 5a-c). AlphaFold 3 modelling indicates that G649 is tightly packed in a hydrophobic pocket (Figure 5d), whilst its backbone dihedral angles fall within the generally favoured region of the Ramachandran plot (Figure 5e). Substitution with a long, charged arginine side-chain would likely cause severe steric clash with the hydrophobic pocket. Hence, similar to P638L, the G649R mutation is predicted to disrupt local MSH4 folding or stability and/or induce a conformational change that impairs function. Using CRISPR-Cas9 genome editing we created a *C. elegans* strain carrying the corresponding *C. elegans* mutation (*msh-4^G557R^)* (Figure 5b) and assessed fertility of worms homozygous for this mutation. *msh-4^G557R^* mutant worms displayed 98% embryonic lethality compared to 0.1% in wild type controls (Figure 5f). This level of embryonic lethality is similar to that observed in *msh-4^P546L^* mutants (modelling human variant *MSH4^P638L^*) and *msh-4* null mutants (Figure 1f-g), suggesting that the G557R substitution impairs MSH-4 function in *C. elegans*. To directly assess the impact of the G557R substitution on meiotic recombination, we next investigated chiasma formation in *msh-4^G557R^* mutants. In contrast to the 6 bivalents present in diakinesis oocytes of control worms, diakinesis oocytes of *msh-4^G557R^* mutants displayed a near complete absence of bivalents (Figure 5g), confirming a severe impairment of meiotic recombination. These results confirm that the G557R substitution results in a *msh-4* null mutant-like phenotype in *C. elegans*, suggesting that the *MSH4^G649R^* variant, when found in homozygosity, is likely to induce infertility in humans.

**Figure 5.**
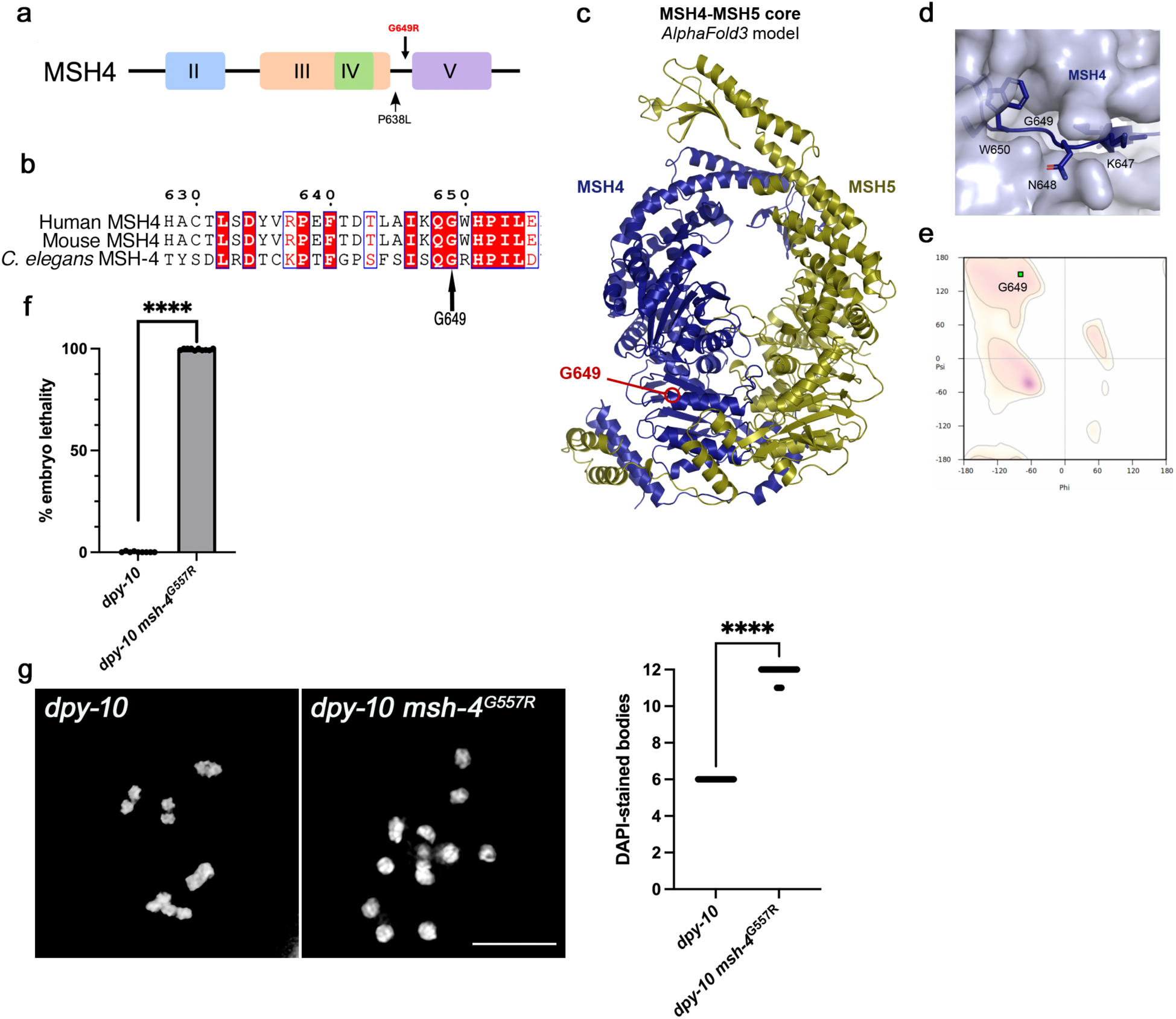
*C. elegans msh-4^G557R^* mutants display viability and recombination defects. **(a)** Diagram showing the location of the human variant G649R in the MSH4 protein. **(b)** Protein sequence alignment showing that G649 in human MSH4 corresponds to G557 in *C. elegans* MSH-4. Numbering refers to the human protein. **(c)** *AlphaFold 3* model of the structure core of a human MSH4-MSH5 1:1 complex (MSH4, amino acids 148-926, blue; MSH5, amino acids 47-834, yellow), highlighting the location of amino acid G649. **(d)** Structural environment of G649 within the model, showing how local side-chains are predicted to pack against the remainder of the MSH4 surface. **(e)** Ramachandran plot showing that the predicted dihedral angles of amino acid G649 are within the favoured region for general amino acids. **(f)** Quantification of embryonic lethality of indicated genotypes, note the increased lethality in *dpy-10 msh-4^G557R^* mutants compared to *dpy-10* controls (the *dpy-10* mutation was introduced during the CRISPR-Cas9 protocol as it is used as a marker for successful injections and therefore was used as the wild-type control for this experiment). Graph shows quantification of the percentage of dead embryos amongst the progeny of worms of indicated genotypes (progeny from 9 worms were scored for *dpy-10* and from 10 worms for *dpy-10 msh-4^G557R^*, error bars indicate mean with 95% CI, p values were calculated using a two-tailed Mann-Whitney U test (**** indicates p < 0.0001). **(g)** Projections of nuclei from diakinesis oocytes stained with DAPI, note the presence of 6 DAPI-stained bodies in the *dpy-10* (wild type) nucleus and of 12 DAPI-stained bodies in the *dpy-10 msh-4^G557R^* mutant nucleus. Graph shows quantification of the number of DAPI-stained bodies in diakinesis oocytes of indicated genotypes (30 nuclei scored per genotype), error bars indicate mean with 95% CI, p values were calculated using a two-tailed Mann-Whitney U test (**** indicates p < 0.0001). Scale bar = 5 µm.

We next modelled in worms the *MSH4^Y589C^* human variant (corresponding to *C. elegans msh-4^F497C^* (Figure 6a)), which has a 0.182 (likely benign) pathogenicity score in AlphaMissense and has a high prevalence in homozygosity in the human population, thus suggesting that it is not causative of infertility. Indeed, AlphaFold 3 modelling places Y589 in a structured but partially solvent-exposed region (Figures 6b-c), in which substitution with a cysteine residue is not predicted to cause major steric clashes, suggesting that the mutation may be structurally tolerated. Using CRISPR-Cas9 genome editing we created a *C. elegans* strain carrying the corresponding *C. elegans* mutation (*msh-4^F497C^)* (Figure 6a). *msh-4^F497C^* mutant worms display levels of embryonic lethality similar to wild type controls, suggesting that meiotic recombination is not impaired by the F497C substitution in MSH-4 (Figure 6d). In agreement with this possibility, chiasma formation is not affected in diakinesis oocytes of *msh-4^F497C^* mutants (Figure 6e). Thus, the F497C substitution does not detectably affect MSH-4 function in *C. elegans*, in agreement with the high number of homozygous cases detected for the corresponding human variant (*MSH4^Y589C^*). These experiments suggest that combining in-silico pathogenicity predictions and structural modelling of human proteins with in-vivo modelling in *C. elegans* provides a powerful and efficient approach for determining the impact of genetic variants found in the human population on fertility.

**Figure 6.**
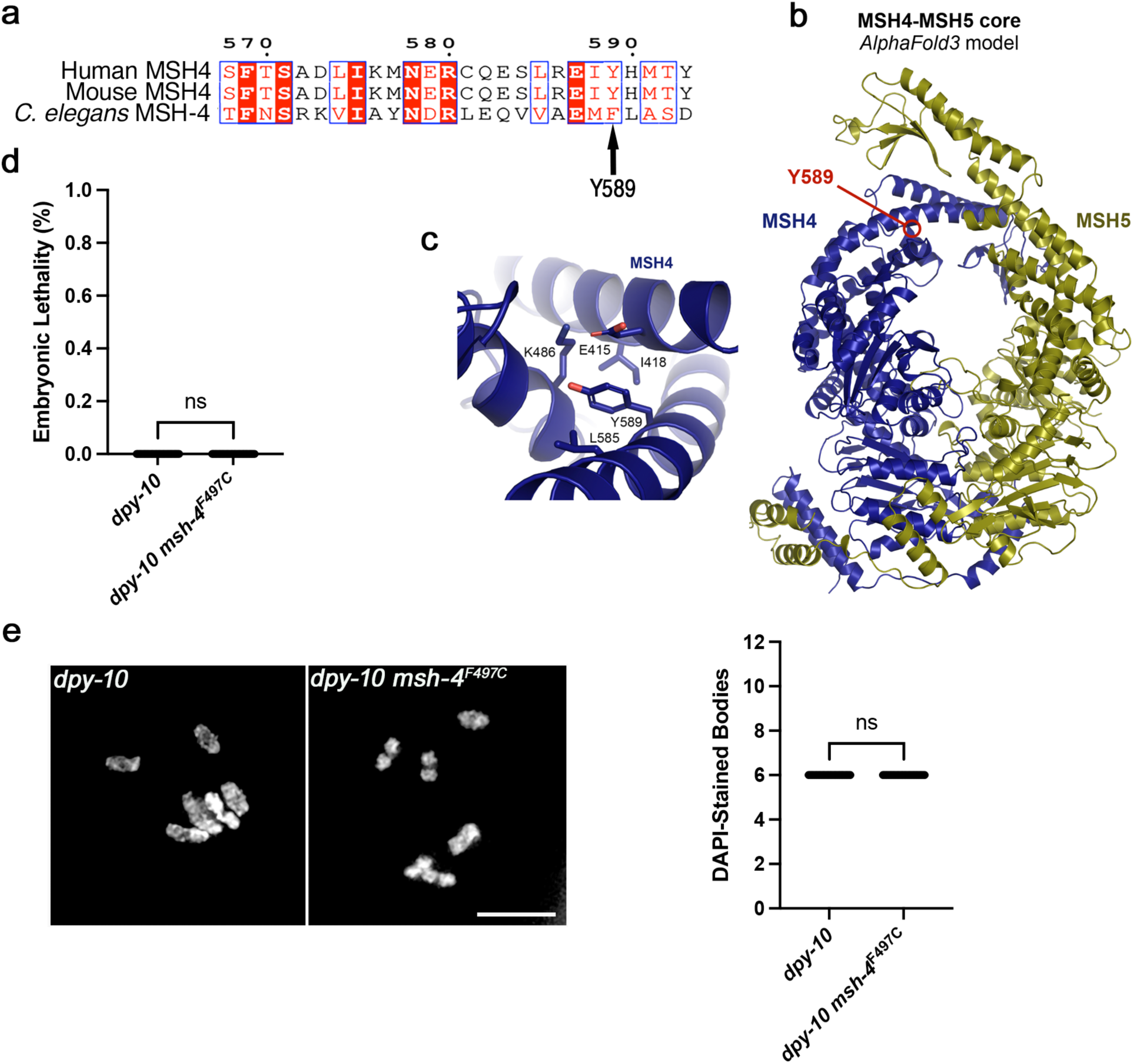
*C. elegans msh-4^F497C^* mutants display normal fertility and recombination. **(a)** Protein sequence alignment showing that Y589 in human MSH4 corresponds to F497 in *C. elegans* MSH-4. Numbering refers to the human protein. **(b)** *AlphaFold 3* model of the structure core of a human MSH4-MSH5 1:1 complex (MSH4, amino acids 148-926, blue; MSH5, amino acids 47-834, yellow), highlighting the location of amino acid Y589. **(c)** Structural environment of Y589 within the model, showing how the tyrosine side-chain is partially solvent-exposed. **(d)** Quantification of embryonic lethality and total number of embryos laid per worm (brood size), note the lack of significant differences between *dpy-10 msh-4^F497C^* mutants and *dpy-10* controls (the *dpy-10* mutation, which reduces body size, was introduced during the CRISPR-Cas9 protocol as it is use as a marker for successful injections and therefore used as the wild-type control for this experiment). 10 worms were scored per genotype, average percentage of dead embryos and average brood size per genotype were compared using the Mann-Whitney U test (“ns” indicates p> 0.05). **(e)** Projections of nuclei from diakinesis oocytes stained with DAPI, note the presence of 6 DAPI-stained bodies in both *dpy-10* (wild type) and *dpy-10 msh-4^F497C^* mutant nucleus. Graph shows quantification of the number of DAPI-stained bodies in indicated genotypes (30 nuclei scored per genotype and compared using the Mann-Whitney U test (“ns” indicates p> 0.05). Scale bar = 5 µm.

## Discussion

The primary aim of this study was to establish a functional pipeline for evaluating genetic variants identified in patients with infertility and elucidating the mechanisms through which these variants impair fertility. The need for this functional pipeline arises from the increasing number of infertility-associated mutations identified by exome and whole genome sequencing of infertile patients. For example, by 2025, publicly available data from sequencing of NOA patients had identified 233 genes and 981 genetic variants potentially causative of infertility ^1^. In the absence of functional validation while genetic variants that induce deletions or premature stop codons in genes previously known to affect fertility may be catalogued as likely pathogenic, assigning pathogenicity to genetic variants encoding missense mutations is much more difficult, even for variants in genes known to play a role in fertility. To address this need and as a proof of concept of our approach, we decided to model the *MSH4^P638L^* variant identified in homozygosis in a NOA patient ^27^. Our worm and mouse models clearly demonstrate that the P638L (P546L in worms and P660L in mouse) substitution perturbs progression of meiotic recombination, with both models displaying phenotypes reminiscent of the corresponding *Msh4* null mutant, including absence of sperm in mouse. These results support that *MSH4^P638L^* causes NOA when present in homozygosity. Moreover, homozygous *Msh4^P660L^* female mice display complete absence of oocyte-containing follicles in their ovaries, demonstrating that this variant also impairs oogenesis. Therefore, we predict that homozygosity for the *MSH4^P638L^* variant will also cause POI in women. Interestingly, the family tree of the *MSH4^P638L^* NOA patient includes three sisters with offspring and two infertile sisters ^27^. Although none of these sisters were genotyped, it is possible that the infertile sisters may also be homozygous for *MSH4^P638L^* and therefore display POI, as previously observed in women homozygous for a splice site mutation in *MSH4* ^32^. Our modelling of *MSH4^P638L^* in worms and mice demonstrates that these organisms can be used to functionally validate potential infertility-inducing genetic variants identified in patients and to elucidate the disease mechanisms affecting these individuals.

Our cytological analysis of spermatogenesis in homozygous *Msh4^P660L^* mice demonstrates complete absence of sperm due to meiotic arrest, as observed in the infertile patient homozygous for *MSH4^P638L^* ^27^. However, *Msh4^P660L^* mice display arrest during early zygotene-like stages of meiotic prophase, as previously observed in *Msh4* null mutants ^18^, while testis sections of the *MSH4^P638L^* patient revealed that at least some cells progress to arrest at metaphase I ^27^, a later meiotic stage. Modelling the P638L substitution in MSH4 in silico, in *C. elegans* and in mouse all suggest that this variant likely severely impairs MSH4 function to a similar extent as a null allele. The difference in the point of meiotic arrest between mice and humans carrying the equivalent MSH4 variant therefore may reflect this variant having species-specific consequences for MSH4 function, or the quality control mechanisms that elicit apoptosis of spermatocytes with synapsis and recombination defects being less robust in humans than in mice. We support the latter as the most likely explanation for the different point of arrest.

Early prophase arrest, as displayed by *Msh4^P660L^* mutants, is observed in many mouse meiotic mutants displaying extensive asynapsis, which hinders meiotic sex chromosome inactivation (MSCI), thereby causing inappropriate expression of Y-encoded *Zfy1* and *Zfy2* genes that promote pachytene arrest and cell death ^13,23,51^. Metaphase I arrest is observed in a subset of mouse meiotic mutants, including *Hei10* ^52^, *Mlh1* ^53^, and *Mlh3* ^54^ mutants, that achieve substantial synapsis and have later defects in maturation of recombination intermediates into inter-homologue CO events. The resulting univalent chromosomes activate the spindle assembly checkpoint to trigger cell death at metaphase I (Burgoyne *et al.*, 2009). Spermatocytes with defects in synapsis or recombination that evade a leaky human pachytene checkpoint are likely to contain univalents that can similarly activate the spindle assembly checkpoint at metaphase I.

Some human NOA patients exhibit asynapsis, MSCI failure and cell death during meiotic prophase, suggesting that some features of the mouse pachytene response to asynapsed chromosome axes may operate in human spermatocytes ^55^.However, the inter-tubule heterogeneity in pachytene or metaphase I arrest reported for some *MSH4* or *MSH5* variant NOA patients ^30^ could reflect human spermatocytes having a less robust pachytene synapsis checkpoint. Furthermore, MSCI itself has been reported to be more variable between and within individuals in humans than in mice ^56^, suggesting that a synapsis checkpoint based on MSCI failure in humans might be less robust. In addition, humans possess only a single Y-encoded paralog of *Zfy1* and *Zfy2*, and differences in the expression and function of the mouse and human paralogs can also contribute to the robustness of any synapsis checkpoint based on MSCI failure ^57^. Our finding that the same severely functionally impaired genetic variant elicits a differential point of arrest in human and mouse spermatogenesis therefore may reflect differences in meiotic checkpoint sensitivity between mice and humans. This has important implications for the interpretation of future mouse models of human fertility-associated mutations.

In addition to modelling the infertility-associated *MSH4^P638L^* variant, we have also created worm models of two *MSH4* variants (G649R and Y589C) present in the human population (gnomAD, ^58^), but that have not previously related to infertility. The goal of these experiments was to determine whether worm models can be used to predict the impact of existing human variants on fertility. The variant encoding the Y589C substitution has been identified in 15201 genotypes, including 123 times in homozygosity, making it one of the most common *MSH4* missense variants. In contrast, the variant encoding the G649R substitution has only been identified in 8 genotypes and never in homozygosity. Worms modelling Y589C displayed normal levels of embryonic viability and CO formation during meiosis, demonstrating normal MSH-4 function. On the other hand, worms modelling G649R displayed severely reduced embryonic viability and lack of COs during meiosis, demonstrating impaired MSH-4 function. These in vivo phenotypes match the structural predictions: P638L and G649R are expected to perturb tightly packed hydrophobic pockets in the MSH4 interdomain region, whereas Y589C should be tolerated because the residue is partially solvent exposed. Hence, structural modelling can help prioritise variants for testing in vivo. This primary screen based on two easily scorable phenotypes shows how worm models can provide a quick and affordable system to functionally screen the effect of human genetic variants on meiosis and therefore on fertility. Models of variants for which a clear phenotype is observed in this primary phenotypic screen, such as for *MSH4^G649R^*, can then be used for more in depth mechanistic analysis aiming to identify the primary defect induced by the genetic variant under study. In addition, this primary worm-based screen can be used to subsequently select genetic variants to be modelled in mouse to obtain functional validation in a mammalian system. This approach will also facilitate focused efforts to build a functional predictive atlas of existing genetic variants in genes that appear frequently associated with human infertility such as *MSH4*.

Beyond the validation of recessive infertility-associated variants, worm models offer several opportunities for expanding the functional interpretation of human infertility genetics. First, given the conservation of meiotic pathways between male and female gametogenesis, *C. elegans* could provide a rapid means of assessing whether variants identified in men with infertility may also impair female fertility, thereby informing their potential reproductive consequences in both sexes. Second, worm models can be used to rapidly test a potential meiotic role of genes identified by spatial transcriptomics of human testis or ovaries that had not previously been connected with fertility. Third, worm models could also be used to investigate more complex modes of inheritance, including compound heterozygosity, a genetic configuration frequently identified by sequencing of infertile patients ^1^, and potential dominant causes of infertility. The latter represents an important opportunity, as dominant infertility can be particularly challenging to establish genetically because of extensive genetic heterogeneity and limited recurrence of candidate variants between affected individuals ^5^, while large numbers of heterozygous missense variants remain of uncertain significance. In summary, modelling in *C. elegans* could enable functional screening of large numbers of candidate genes and variants, providing a scalable approach to identify and functionally characterise both recessive and dominant genetic causes of human infertility.

## Materials and Methods

### C. elegans Methods

#### C. elegans genetics

*C. elegans* strains were cultured on NG agar plates containing OP-50 *E. coli* as a source of food and maintained at 20°C. The N2 Bristol strain was used as a wild type control. Strains used in this study are listed in Table S1.

#### CRISPR/Cas9 genome editing

To generate transgenic *C. elegans* lines, L4-stage N2 worms were picked the day prior to microinjection to synchronise their age for injection as young adults. CRISPR-Cas9 mediated genome editing was performed using Cas9 ribonucleoprotein complexes and single-stranded DNA (ssDNA) repair templates, using co-editing of the *dpy-10* locus as a marker to identify edited worms ^59^. CRISPR reagents are shown in Table S2. The injection mix contained the following reagents: 2µl of 100µM target crRNA:trRNA duplex, 2.95µl of Cas9 (at 61 µM), 0.2µl of *dpy-10* crRNA:trRNA duplex, 0.6µl ssDNA repair template (100µM target ssDNA), 0.5µl *dpy-10* ssDNA repair template (10µM *dpy-10* ssDNA) and 3.75µl water. Halocarbon Oil (Sigma) was used to fix the worms onto dried 2% agarose pads, and microneedles were pulled using a micropipette puller (Sutter) and subsequently used to inject worm gonads with 0.4µl of the above injection mixture using the Narishige IM-21 microinjector attached to an Olympus IX71 microscope.

Worms were recovered at 25°C for 4 days, and plates were checked for the presence of the co-CRISPR genotype (dumpy in homozygotes and roller in heterozygotes). Dumpy, roller and wild type worms from positive plates were picked onto individual plates and left for 48 hours to lay eggs. F1 progenies were placed in a PCR tube containing 15µl lysis buffer with Proteinase K. Tubes were microfuged for 15 seconds, then frozen at -80°C for 10 minutes. The tubes were lysed under standard conditions to release DNA (heat to 65°C for 90 minutes then heat to 95°C for 15 minutes). Oligonucleotides were designed to flank the region of interest and to produce a gel electrophoresis band in the N2 strain as a negative control following PCR. PCR products were subsequently digested using a restriction endonuclease (NEB) to detect the presence of the mutation of interest. Homozygous progenies were detected and validated by Sanger sequencing. Heterozygous F2 progeny were outcrossed with wild type male worms and subsequently crossed with a genetic balancer containing a fluorescent marker. A fluorescence microscope (Leica MZ16F) was used to pick worms that were heterozygous for the fluorescent marker to maintain strains. Table S2 provides a list of oligos used to create and genotype CRISPR edits in the *C. elegans* genome.

#### Embryonic lethality

L4-stage worms from each strain were individually picked onto single plates and transferred onto fresh plates every 24 hours until no offspring were produced. Numbers of live progeny and unhatched eggs were counted at 24 hours after worm transfer as viable embryos hatch within 14 hours of being laid, while dead embryos remain unhatched. N2 worms were used as a control strain during lethality experiments. Lethality for each worm was determined as the proportion of unhatched eggs divided by the total number of embryos laid.

#### Immunofluorescence sample preparation and microscopy

Worms were dissected in 1x Egg Buffer (118 mM NaCl, 48 mM KCl_2_, 2 mM CaCl_2_, 2 mM MgCl_2_, 25 mM HEPES pH7.4, 0.1% Tween20) and were fixed in 1x Egg Buffer with 1% paraformaldehyde for five minutes on a SuperFrost Plus slide (VWR). Slides were then flash frozen in liquid nitrogen and the cover slip was removed before the slides were placed in ice cold methanol for 1 minute. Slides were washed in 1xPBST (1x PBS, 0.1% Tween20) 3x for ten minutes before they were placed in Block (1xPBST with 0.7% bovine serum albumin (Sigma-Aldrich, #SLCB3716)) for a minimum of one hour. 50μL of primary antibodies (rabbit α-RAD-51 (1:3000, Nicola Silva) or rabbit HA-AbTag (1:200, Cell Signalling, #2367S)) diluted in PBST were added to each slide and samples were incubated for 16-18 hours in a dark humidifying chamber with parafilm coverslips. Slides were then washed 3x in 1xPBST for 10 minutes. 50μL of secondary antibody (Alexa Fluor 555 Goat α-Rabbit (Invitrogen A21428) or Alexa Fluor 488 Goat α-Rabbit (Invitrogen A11008) diluted 1:500 in PBST were then added to each sample and slides were incubated for two hours in a dark humidifying chamber with parafilm coverslips. Slides were washed 3x in 1xPBST for ten minutes, and then 50μL of 2μg/mL DAPI was applied to each slide. Samples were incubated in a dark humidifying chamber with parafilm coverslips for five minutes, then washed 1x in PBST for five minutes. Slides were mounted in Vectashield mounting media (Vector laboratories) with a coverslip (VWR) and sealed with nail polish. All slides were maintained at 4°C until imaging.

Immunofluorescence images were acquired at 512x512 pixel dimensions on an Applied Precision DeltaVision microscope with a 100x lens and a 1.5x optivar. All images were acquired in 3 dimensions with Z-stacks at 0.2μm intervals. Images were deconvolved with Applied Precision softWoRx software and individual image tiles were stitched into a single image for analysis using the Grid/Collection Stitching module in Adobe Photoshop.

#### DAPI body quantification

Worms were stained with 4’6-diamidino-2-phenylindole (DAPI) using the same protocol as above and the number of DAPI-stained bodies in the nuclei of -1 or -2 diakinesis oocytes of different genotypes were counted on the microscope.

#### Image analysis and quantification

RAD-51 and COSA-1::HA foci were quantified manually using Photoshop. All statistics were calculated in Graphpad Prism (v10.4.2). Comparisons between two groups were performed using a Mann-Whitney U Test, and p values were taken to be significant if < 0.05.

## Mouse Methods

### CRISPR/Cas9 genome editing

Msh4^P660L^ and Msh4^PAM^ mouse lines were generated using CRISPR/Cas9 gene editing, following a similar protocol as previously detailed ^60,61^. gRNA complexes were prepared using Msh4guide crRNA (Table S3) and tracrRNA (IDT), and microinjected at 20ng/μl along with 20ng/ul Msh4_P660L or Msh4_PAM repair oligo and 20ng/μl Alt-R S.p. Cas9 Nuclease V3 (IDT) into zygotes from B6CBAF1 mice. Genotyping of G_0_ mice and their offspring was carried out from ear clip samples, both by restriction digests of the PCR product obtained using primers Msh4_2L and Msh4_2R (Table S3) and by sequencing of PCR products from the same primers. Founder mice for each line were backcrossed to C57BL/6J (Charles River) and the resulting heterozygous mice crossed to generate both wild-type and homozygous animals for phenotyping.

### Mouse phenotyping

Adult mice were culled at 1.5-4 months via cervical dislocation, and either the testes and epididymides or the ovaries dissected in PBS. Testis weights and caudal epididymal sperm counts were obtained as described previously ^62^. For testis and ovary histology, tissues were fixed overnight in Bouin’s fixative at 4°C, then embedded in wax, sectioned and stained with haematoxylin and eosin as previously detailed ^62^. Effects of genotype on testis weights and sperm counts were assessed using Student’s t-tests in R ^63^.

### Meiotic chromosome spreads

Preparation of meiotic chromosome spreads from adult mouse testes was carried out as previously described ^64^. Antibody staining and imaging of the spreads was also performed as previously described ^65^. Primary antibodies were mouse anti-SYCP3 (AbCam, #97672, 1:500), rabbit anti-SYCP3 (LSBio, #LS-B175, 1:500), rabbit anti-SYCP1 (Abcam, #15090, 1:200), mouse anti-γH2AX (Millipore, #05-636, 1:2000), rabbit anti-RAD51 (Millipore, #PC130, 1:500) and mouse anti-MLH1 (BD-Pharmigen, #51-1327GR, 1:50). Slides were mounted using Vectashield anti-fade mounting media (Vector laboratories, H-1200) and high-precision coverslips (Deckgläser, ES0107222).

### Widefield imaging & image analysis

Chromosome spread imaging was performed using a Zeiss Axioimager 2. Image analysis was performed using ImageJ, with counts of RAD51 foci obtained using a custom macro previously designed for image segmentation and meiotic chromosome analysis ^66^. For chromosome spread imaging, staging of meiotic nuclei was based on immunostaining for SYCP3 along chromosome axes. Nuclei with only short, unsynapsed axis segments were defined as leptotene; those with incomplete axis segments with partial synapsis as zygotene; those with fully extended axes with complete synapsis of autosomal chromosomes as pachytene; and those with pairs of extended axes containing regions of desynapsis as diplotene. Zygotene-like nuclei were defined as containing fully extended but completely asynapsed axes or containing at least one completely synapsed pair of autosomes and at least one incompletely synapsed pair of autosomes exhibiting asynapsis along at least half its length ^65^. Nuclei were classified as MLH1-positive if they contained at least fifteen axis-associated MLH1 foci. Effects of genotype on nominal and binary categorical data were analysed by fitting a Bayesian categorical generalised linear mixed model (GLMM) using the *brms* package in R with a logit-linked categorical or logit-linked binomial response distribution respectively ^67^. Effects of genotype on count data were similarly analysed by fitting a Bayesian GLMM using *brms* and a log-linked negative binomial response distribution. Individual mice were treated as random-effect intercepts to prevent pseudoreplication, and datasets with complete separation of zero-variance categories were accommodated by incorporating weakly informative normal priors. Directional Bayesian hypothesis testing was used to determine the posterior probabilities (PPs); a PP > 0.95 was considered statistically significant.

### Structural modelling

Models of the human MSH4-MSH5 complex were generated by *AlphaFold 3* using the *AlphaFold* Server ^68^. The pLDDT and PAE plots were generated using https://github.com/Ash100/Biopython/blob/main/AF3_Results_Visualization.ipynb. Initially, full-length sequences were used to generate a model of a 1:1 complex. On this basis, unstructured N- and C-terminal sequences were removed, and modelling was repeating using the structural cores of human MSH4 (amino acids 148-926) and MSH5 (amino acids 47-834). Dihedral angles were viewed on Ramachandran plots using *Coot* ^69^. Molecular structure images were generated using the *PyMOL* Molecular Graphics System, Version 3.0 Schrödinger, LLC.

## Acknowledgements

We acknowledge Margot Wyrwoll (University of Edinburgh) for providing critical comments on the manuscript. This work was supported by a MRC Clinical Research Training Fellowship to O.E.T., by a MRC core-funded grant (MC-A652-5PY60) to E.M.-P., and a MRC University Unit award (MC_UU_00035/3) to I.R.A.. O.R.D. is a Wellcome Senior Research Fellow (Grant Number 219413/Z/19/Z).

## References

1. Ibrahim, R., Malcher, A., and Kurpisz, M. (2025). Summarizing the human genes and their variants causative of non-obstructive azoospermia uncovered using whole genome/exome sequencing. Reprod Biol 25, 101048. 10.1016/j.repbio.2025.101048.

2. Touraine, P., Chabbert-Buffet, N., Plu-Bureau, G., Duranteau, L., Sinclair, A.H., and Tucker, E.J. (2024). Premature ovarian insufficiency. Nat Rev Dis Primers 10, 63. 10.1038/s41572-024-00547-5.

3. Krausz, C., and Riera-Escamilla, A. (2018). Genetics of male infertility. Nat Rev Urol 15, 369–384. 10.1038/s41585-018-0003-3.

4. Houston, B.J., Riera-Escamilla, A., Wyrwoll, M.J., Salas-Huetos, A., Xavier, M.J., Nagirnaja, L., Friedrich, C., Conrad, D.F., Aston, K.I., Krausz, C., et al. (2021). A systematic review of the validated monogenic causes of human male infertility: 2020 update and a discussion of emerging gene-disease relationships. Hum Reprod Update 28, 15–29. 10.1093/humupd/dmab030.

5. Oud, M.S., Smits, R.M., Smith, H.E., Mastrorosa, F.K., Holt, G.S., Houston, B.J., de Vries, P.F., Alobaidi, B.K.S., Batty, L.E., Ismail, H., et al. (2022). A de novo paradigm for male infertility. Nat Commun 13, 154. 10.1038/s41467-021-27132-8.

6. Nagirnaja, L., Morup, N., Nielsen, J.E., Stakaitis, R., Golubickaite, I., Oud, M.S., Winge, S.B., Carvalho, F., Aston, K.I., Khani, F., et al. (2021). Variant PNLDC1, Defective piRNA Processing, and Azoospermia. N Engl J Med 385, 707–719. 10.1056/NEJMoa2028973.

7. Riera-Escamilla, A., Vockel, M., Nagirnaja, L., Xavier, M.J., Carbonell, A., Moreno-Mendoza, D., Pybus, M., Farnetani, G., Rosta, V., Cioppi, F., et al. (2022). Large-scale analyses of the X chromosome in 2,354 infertile men discover recurrently affected genes associated with spermatogenic failure. Am J Hum Genet 109, 1458–1471. 10.1016/j.ajhg.2022.06.007.

8. Adams, I.R., and Davies, O.R. (2023). Meiotic Chromosome Structure, the Synaptonemal Complex, and Infertility. Annu Rev Genomics Hum Genet 24, 35–61. 10.1146/annurev-genom-110122-090239.

9. Saitou, M., Nagano, M., and Mizuta, K. (2026). Mechanisms of human germ cell development. Nat Rev Mol Cell Biol 27, 153–171. 10.1038/s41580-025-00893-6.

10. Petronczki, M., Siomos, M.F., and Nasmyth, K. (2003). Un menage a quatre: the molecular biology of chromosome segregation in meiosis. Cell 112, 423–440. 10.1016/s0092-8674(03)00083-7.

11. Grey, C., and de Massy, B. (2021). Chromosome Organization in Early Meiotic Prophase. Front Cell Dev Biol 9, 688878. 10.3389/fcell.2021.688878.

12. Hunter, N. (2015). Meiotic Recombination: The Essence of Heredity. Cold Spring Harb Perspect Biol 7. 10.1101/cshperspect.a016618.

13. Burgoyne, P.S., Mahadevaiah, S.K., and Turner, J.M. (2009). The consequences of asynapsis for mammalian meiosis. Nat Rev Genet 10, 207–216. 10.1038/nrg2505.

14. Romanienko, P.J., and Camerini-Otero, R.D. (2000). The mouse Spo11 gene is required for meiotic chromosome synapsis. Mol Cell 6, 975–987. 10.1016/s1097-2765(00)00097-6.

15. de Vries, F.A., de Boer, E., van den Bosch, M., Baarends, W.M., Ooms, M., Yuan, L., Liu, J.G., van Zeeland, A.A., Heyting, C., and Pastink, A. (2005). Mouse Sycp1 functions in synaptonemal complex assembly, meiotic recombination, and XY body formation. Genes Dev 19, 1376–1389. 10.1101/gad.329705.

16. Wang, X., Nie, L., Hong, Z., Li, L., Fan, Q., Ma, B., Li, Z., Gao, Y., Zhang, M., Zhang, Y., and Wang, M. (2025). Genetic and epigenetic insights into non-obstructive azoospermia: mechanisms, biomarkers, and clinical perspectives. Reprod Biol Endocrinol 23, 159. 10.1186/s12958-025-01497-7.

17. Ross-Macdonald, P., and Roeder, G.S. (1994). Mutation of a meiosis-specific MutS homolog decreases crossing over but not mismatch correction. Cell 79, 1069–1080. 10.1016/0092-8674(94)90037-x.

18. Kneitz, B., Cohen, P.E., Avdievich, E., Zhu, L., Kane, M.F., Hou, H., Jr., Kolodner, R.D., Kucherlapati, R., Pollard, J.W., and Edelmann, W. (2000). MutS homolog 4 localization to meiotic chromosomes is required for chromosome pairing during meiosis in male and female mice. Genes Dev 14, 1085–1097.

19. Zalevsky, J., MacQueen, A.J., Duffy, J.B., Kemphues, K.J., and Villeneuve, A.M. (1999). Crossing over during Caenorhabditis elegans meiosis requires a conserved MutS-based pathway that is partially dispensable in budding yeast. Genetics 153, 1271–1283. 10.1093/genetics/153.3.1271.

20. Pochart, P., Woltering, D., and Hollingsworth, N.M. (1997). Conserved properties between functionally distinct MutS homologs in yeast. J Biol Chem 272, 30345–30349. 10.1074/jbc.272.48.30345.

21. Cannavo, E., Sanchez, A., Anand, R., Ranjha, L., Hugener, J., Adam, C., Acharya, A., Weyland, N., Aran-Guiu, X., Charbonnier, J.B., et al. (2020). Regulation of the MLH1-MLH3 endonuclease in meiosis. Nature 586, 618–622. 10.1038/s41586-020-2592-2.

22. Snowden, T., Acharya, S., Butz, C., Berardini, M., and Fishel, R. (2004). hMSH4-hMSH5 recognizes Holliday Junctions and forms a meiosis-specific sliding clamp that embraces homologous chromosomes. Mol Cell 15, 437–451. 10.1016/j.molcel.2004.06.040.

23. Mahadevaiah, S.K., Bourc’his, D., de Rooij, D.G., Bestor, T.H., Turner, J.M., and Burgoyne, P.S. (2008). Extensive meiotic asynapsis in mice antagonises meiotic silencing of unsynapsed chromatin and consequently disrupts meiotic sex chromosome inactivation. J Cell Biol 182, 263–276. 10.1083/jcb.200710195.

24. de Vries, S.S., Baart, E.B., Dekker, M., Siezen, A., de Rooij, D.G., de Boer, P., and te Riele, H. (1999). Mouse MutS-like protein Msh5 is required for proper chromosome synapsis in male and female meiosis. Genes Dev 13, 523–531. 10.1101/gad.13.5.523.

25. Edelmann, W., Cohen, P.E., Kneitz, B., Winand, N., Lia, M., Heyer, J., Kolodner, R., Pollard, J.W., and Kucherlapati, R. (1999). Mammalian MutS homologue 5 is required for chromosome pairing in meiosis. Nat Genet 21, 123–127. 10.1038/5075.

26. Milano, C.R., Holloway, J.K., Zhang, Y., Jin, B., Smith, C., Bergman, A., Edelmann, W., and Cohen, P.E. (2019). Mutation of the ATPase Domain of MutS Homolog-5 (MSH5) Reveals a Requirement for a Functional MutSgamma Complex for All Crossovers in Mammalian Meiosis. G3 (Bethesda) 9, 1839–1850. 10.1534/g3.119.400074.

27. Krausz, C., Riera-Escamilla, A., Moreno-Mendoza, D., Holleman, K., Cioppi, F., Algaba, F., Pybus, M., Friedrich, C., Wyrwoll, M.J., Casamonti, E., et al. (2020). Genetic dissection of spermatogenic arrest through exome analysis: clinical implications for the management of azoospermic men. Genet Med 22, 1956–1966. 10.1038/s41436-020-0907-1.

28. Akbari, A., Padidar, K., Salehi, N., Mashayekhi, M., Almadani, N., Sadighi Gilani, M.A., Bashambou, A., McElreavey, K., and Totonchi, M. (2021). Rare missense variant in MSH4 associated with primary gonadal failure in both 46, XX and 46, XY individuals. Hum Reprod 36, 1134–1145. 10.1093/humrep/deaa362.

29. Li, P., Ji, Z., Zhi, E., Zhang, Y., Han, S., Zhao, L., Tian, R., Chen, H., Huang, Y., Zhang, J., et al. (2022). Novel bi-allelic MSH4 variants causes meiotic arrest and non-obstructive azoospermia. Reprod Biol Endocrinol 20, 21. 10.1186/s12958-022-00900-x.

30. Wyrwoll, M.J., van Walree, E.S., Hamer, G., Rotte, N., Motazacker, M.M., Meijers-Heijboer, H., Alders, M., Meissner, A., Kaminsky, E., Woste, M., et al. (2021). Bi-allelic variants in DNA mismatch repair proteins MutS Homolog MSH4 and MSH5 cause infertility in both sexes. Hum Reprod 37, 178–189. 10.1093/humrep/deab230.

31. Hashemi Sheikhshabani, S., Ghafouri-Fard, S., Hosseini, E., and Omrani, M.D. (2024). A novel homozygote nonsense variant of MSH4 leads to primary ovarian insufficiency and non-obstructive azoospermia. Mol Biol Rep 51, 68. 10.1007/s11033-023-09000-4.

32. Carlosama, C., Elzaiat, M., Patino, L.C., Mateus, H.E., Veitia, R.A., and Laissue, P. (2017). A homozygous donor splice-site mutation in the meiotic gene MSH4 causes primary ovarian insufficiency. Hum Mol Genet 26, 3161–3166. 10.1093/hmg/ddx199.

33. Dernburg, A.F., McDonald, K., Moulder, G., Barstead, R., Dresser, M., and Villeneuve, A.M. (1998). Meiotic recombination in C. elegans initiates by a conserved mechanism and is dispensable for homologous chromosome synapsis. Cell 94, 387–398. 10.1016/s0092-8674(00)81481-6.

34. Alpi, A., Pasierbek, P., Gartner, A., and Loidl, J. (2003). Genetic and cytological characterization of the recombination protein RAD-51 in Caenorhabditis elegans. Chromosoma 112, 6–16. 10.1007/s00412-003-0237-5.

35. Hinch, A.G., Becker, P.W., Li, T., Moralli, D., Zhang, G., Bycroft, C., Green, C., Keeney, S., Shi, Q., Davies, B., and Donnelly, P. (2020). The Configuration of RPA, RAD51, and DMC1 Binding in Meiosis Reveals the Nature of Critical Recombination Intermediates. Mol Cell 79, 689–701 e610. 10.1016/j.molcel.2020.06.015.

36. Yokoo, R., Zawadzki, K.A., Nabeshima, K., Drake, M., Arur, S., and Villeneuve, A.M. (2012). COSA-1 reveals robust homeostasis and separable licensing and reinforcement steps governing meiotic crossovers. Cell 149, 75–87. 10.1016/j.cell.2012.01.052.

37. Colaiacovo, M.P., MacQueen, A.J., Martinez-Perez, E., McDonald, K., Adamo, A., La Volpe, A., and Villeneuve, A.M. (2003). Synaptonemal complex assembly in C. elegans is dispensable for loading strand-exchange proteins but critical for proper completion of recombination. Dev Cell 5, 463–474. 10.1016/s1534-5807(03)00232-6.

38. Zetka, M.C., Kawasaki, I., Strome, S., and Muller, F. (1999). Synapsis and chiasma formation in Caenorhabditis elegans require HIM-3, a meiotic chromosome core component that functions in chromosome segregation. Genes Dev 13, 2258–2270. 10.1101/gad.13.17.2258.

39. MacQueen, A.J., Colaiacovo, M.P., McDonald, K., and Villeneuve, A.M. (2002). Synapsis-dependent and -independent mechanisms stabilize homolog pairing during meiotic prophase in C. elegans. Genes Dev 16, 2428–2442. 10.1101/gad.1011602.

40. Yuan, L., Liu, J.G., Zhao, J., Brundell, E., Daneholt, B., and Hoog, C. (2000). The murine SCP3 gene is required for synaptonemal complex assembly, chromosome synapsis, and male fertility. Mol Cell 5, 73–83. 10.1016/s1097-2765(00)80404-9.

41. Abe, H., Alavattam, K.G., Hu, Y.C., Pang, Q., Andreassen, P.R., Hegde, R.S., and Namekawa, S.H. (2020). The Initiation of Meiotic Sex Chromosome Inactivation Sequesters DNA Damage Signaling from Autosomes in Mouse Spermatogenesis. Curr Biol 30, 408–420 e405. 10.1016/j.cub.2019.11.064.

42. Fernandez-Capetillo, O., Mahadevaiah, S.K., Celeste, A., Romanienko, P.J., Camerini-Otero, R.D., Bonner, W.M., Manova, K., Burgoyne, P., and Nussenzweig, A. (2003). H2AX is required for chromatin remodeling and inactivation of sex chromosomes in male mouse meiosis. Dev Cell 4, 497–508. 10.1016/s1534-5807(03)00093-5.

43. He, W., Rao, H., Tang, S., Bhagwat, N., Kulkarni, D.S., Ma, Y., Chang, M.A.W., Hall, C., Bragg, J.W., Manasca, H.S., et al. (2020). Regulated Proteolysis of MutSgamma Controls Meiotic Crossing Over. Mol Cell 78, 168–183 e165. 10.1016/j.molcel.2020.02.001.

44. Moens, P.B., Kolas, N.K., Tarsounas, M., Marcon, E., Cohen, P.E., and Spyropoulos, B. (2002). The time course and chromosomal localization of recombination-related proteins at meiosis in the mouse are compatible with models that can resolve the early DNA-DNA interactions without reciprocal recombination. J Cell Sci 115, 1611–1622. 10.1242/jcs.115.8.1611.

45. Kauppi, L., Barchi, M., Lange, J., Baudat, F., Jasin, M., and Keeney, S. (2013). Numerical constraints and feedback control of double-strand breaks in mouse meiosis. Genes Dev 27, 873–886. 10.1101/gad.213652.113.

46. Anderson, L.K., Reeves, A., Webb, L.M., and Ashley, T. (1999). Distribution of crossing over on mouse synaptonemal complexes using immunofluorescent localization of MLH1 protein. Genetics 151, 1569–1579. 10.1093/genetics/151.4.1569.

47. Falque, M., Mercier, R., Mezard, C., de Vienne, D., and Martin, O.C. (2007). Patterns of recombination and MLH1 foci density along mouse chromosomes: modeling effects of interference and obligate chiasma. Genetics 176, 1453–1467. 10.1534/genetics.106.070235.

48. Kolas, N.K., Svetlanov, A., Lenzi, M.L., Macaluso, F.P., Lipkin, S.M., Liskay, R.M., Greally, J., Edelmann, W., and Cohen, P.E. (2005). Localization of MMR proteins on meiotic chromosomes in mice indicates distinct functions during prophase I. J Cell Biol 171, 447–458. 10.1083/jcb.200506170.

49. Chen, S., Francioli, L.C., Goodrich, J.K., Collins, R.L., Kanai, M., Wang, Q., Alfoldi, J., Watts, N.A., Vittal, C., Gauthier, L.D., et al. (2024). A genomic mutational constraint map using variation in 76,156 human genomes. Nature 625, 92–100. 10.1038/s41586-023-06045-0.

50. Cheng, J., Novati, G., Pan, J., Bycroft, C., Zemgulyte, A., Applebaum, T., Pritzel, A., Wong, L.H., Zielinski, M., Sargeant, T., et al. (2023). Accurate proteome-wide missense variant effect prediction with AlphaMissense. Science 381, eadg7492. 10.1126/science.adg7492.

51. Royo, H., Polikiewicz, G., Mahadevaiah, S.K., Prosser, H., Mitchell, M., Bradley, A., de Rooij, D.G., Burgoyne, P.S., and Turner, J.M. (2010). Evidence that meiotic sex chromosome inactivation is essential for male fertility. Curr Biol 20, 2117–2123. 10.1016/j.cub.2010.11.010.

52. Ward, J.O., Reinholdt, L.G., Motley, W.W., Niswander, L.M., Deacon, D.C., Griffin, L.B., Langlais, K.K., Backus, V.L., Schimenti, K.J., O’Brien, M.J., et al. (2007). Mutation in mouse hei10, an e3 ubiquitin ligase, disrupts meiotic crossing over. PLoS Genet 3, e139. 10.1371/journal.pgen.0030139.

53. Baker, S.M., Plug, A.W., Prolla, T.A., Bronner, C.E., Harris, A.C., Yao, X., Christie, D.M., Monell, C., Arnheim, N., Bradley, A., et al. (1996). Involvement of mouse Mlh1 in DNA mismatch repair and meiotic crossing over. Nat Genet 13, 336–342. 10.1038/ng0796-336.

54. Lipkin, S.M., Moens, P.B., Wang, V., Lenzi, M., Shanmugarajah, D., Gilgeous, A., Thomas, J., Cheng, J., Touchman, J.W., Green, E.D., et al. (2002). Meiotic arrest and aneuploidy in MLH3-deficient mice. Nat Genet 31, 385–390. 10.1038/ng931.

55. Jan, S.Z., Jongejan, A., Korver, C.M., van Daalen, S.K.M., van Pelt, A.M.M., Repping, S., and Hamer, G. (2018). Distinct prophase arrest mechanisms in human male meiosis. Development 145. 10.1242/dev.160614.

56. de Vries, M., Vosters, S., Merkx, G., D’Hauwers, K., Wansink, D.G., Ramos, L., and de Boer, P. (2012). Human male meiotic sex chromosome inactivation. PLoS One 7, e31485. 10.1371/journal.pone.0031485.

57. Decarpentrie, F., Vernet, N., Mahadevaiah, S.K., Longepied, G., Streichemberger, E., Aknin-Seifer, I., Ojarikre, O.A., Burgoyne, P.S., Metzler-Guillemain, C., and Mitchell, M.J. (2012). Human and mouse ZFY genes produce a conserved testis-specific transcript encoding a zinc finger protein with a short acidic domain and modified transactivation potential. Hum Mol Genet 21, 2631–2645. 10.1093/hmg/dds088.

58. Karczewski, K.J., Francioli, L.C., Tiao, G., Cummings, B.B., Alfoldi, J., Wang, Q., Collins, R.L., Laricchia, K.M., Ganna, A., Birnbaum, D.P., et al. (2020). The mutational constraint spectrum quantified from variation in 141,456 humans. Nature 581, 434–443. 10.1038/s41586-020-2308-7.

59. Paix, A., Schmidt, H., and Seydoux, G. (2016). Cas9-assisted recombineering in C. elegans: genome editing using in vivo assembly of linear DNAs. Nucleic Acids Res 44, e128. 10.1093/nar/gkw502.

60. Quadros, R.M., Miura, H., Harms, D.W., Akatsuka, H., Sato, T., Aida, T., Redder, R., Richardson, G.P., Inagaki, Y., Sakai, D., et al. (2017). Easi-CRISPR: a robust method for one-step generation of mice carrying conditional and insertion alleles using long ssDNA donors and CRISPR ribonucleoproteins. Genome Biol 18, 92. 10.1186/s13059-017-1220-4.

61. Crichton, J.H., Dunce, J.M., Dunne, O.M., Salmon, L.J., Devenney, P.S., Lawson, J., Adams, I.R., and Davies, O.R. (2023). Structural maturation of SYCP1-mediated meiotic chromosome synapsis by SYCE3. Nat Struct Mol Biol 30, 188–199. 10.1038/s41594-022-00909-1.

62. Ollinger, R., Childs, A.J., Burgess, H.M., Speed, R.M., Lundegaard, P.R., Reynolds, N., Gray, N.K., Cooke, H.J., and Adams, I.R. (2008). Deletion of the pluripotency-associated Tex19.1 gene causes activation of endogenous retroviruses and defective spermatogenesis in mice. PLoS Genet 4, e1000199. 10.1371/journal.pgen.1000199.

63. Team, R.C. (2026). R: A Language and Environment for Statistical Computing

64. Costa, Y., Speed, R., Ollinger, R., Alsheimer, M., Semple, C.A., Gautier, P., Maratou, K., Novak, I., Hoog, C., Benavente, R., and Cooke, H.J. (2005). Two novel proteins recruited by synaptonemal complex protein 1 (SYCP1) are at the centre of meiosis. J Cell Sci 118, 2755–2762. 10.1242/jcs.02402.

65. Crichton, J.H., Playfoot, C.J., MacLennan, M., Read, D., Cooke, H.J., and Adams, I.R. (2017). Tex19.1 promotes Spo11-dependent meiotic recombination in mouse spermatocytes. PLoS Genet 13, e1006904. 10.1371/journal.pgen.1006904.

66. Crichton, J.H., and Adams, I.R. (2024). Computational Tools for the Analysis of Meiotic Prophase I Images. Methods Mol Biol 2818, 249–270. 10.1007/978-1-0716-3906-1_17.

67. Bürkner, P.C. (2017). brms: An R Package for Bayesian Multilevel Models Using Stan. J Stat Softw 80, 1–28. 10.18637/jss.v080.i01.

68. Abramson, J., Adler, J., Dunger, J., Evans, R., Green, T., Pritzel, A., Ronneberger, O., Willmore, L., Ballard, A.J., Bambrick, J., et al. (2024). Accurate structure prediction of biomolecular interactions with AlphaFold 3. Nature 630, 493–500. 10.1038/s41586-024-07487-w.

69. Emsley, P., Lohkamp, B., Scott, W.G., and Cowtan, K. (2010). Features and development of Coot. Acta Crystallogr D Biol Crystallogr 66, 486–501. 10.1107/S0907444910007493.

