## Supplementary material for "Functional modelling of human infertility-associated genetic variants using *C. elegans* and mice": Figure S1

### Supplementary Figures

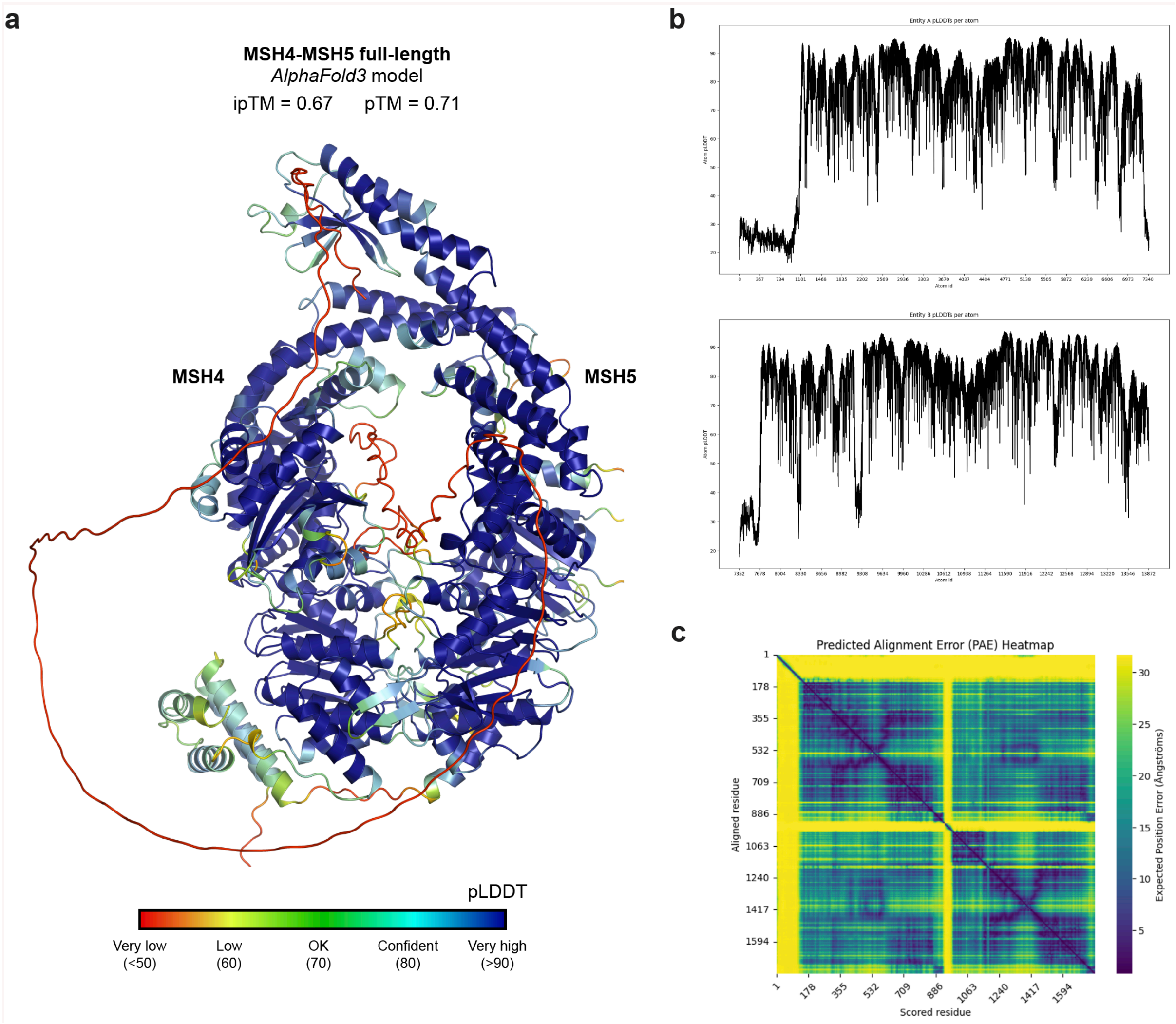


**Figure S1. (a)** *AlphaFold 3* model of the full-length human MSH4-MSH5 1:1 complex coloured according to predicted LDDT (pLDDT) scores, between blue (>90) and red (<50). (**b**) Predicted LDDT (pLDDT) scores shown for each amino acid of the MSH4 chain (top) and MSH5 chain (bottom). (**c**) Predicted aligned error scores between each amino acid of the MSH4 and MSH5 chains, between blue (low error) and yellow (high error).

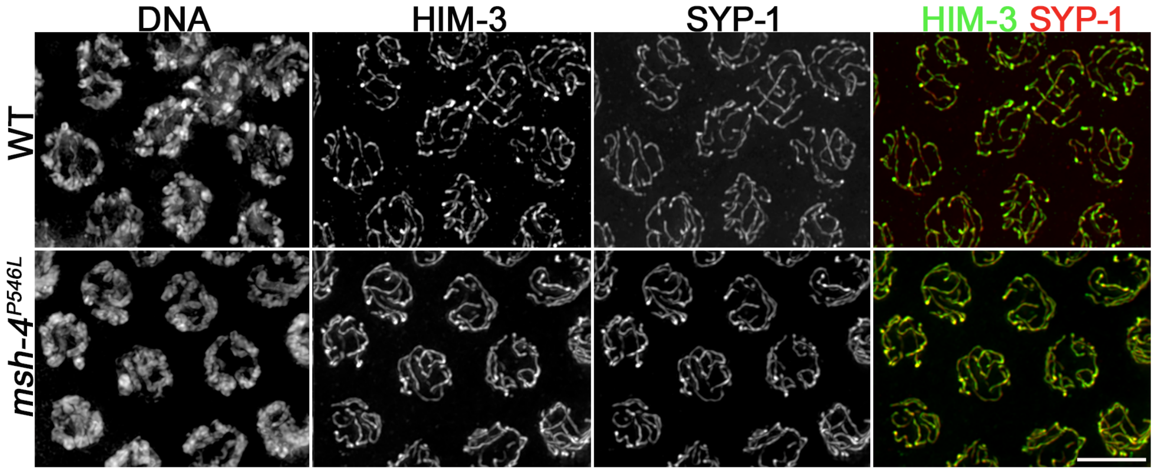


**Figure S2**. Projections of mid pachytene nuclei stained with anti-HIM-3 and anti-SYP-1 antibodies and DAPI. Note that both wild type and *msh-4*$^{P546L}$ mutant nuclei display full overlap of HIM-3 and SYP-1 signals. Scale bar = 5 µm.

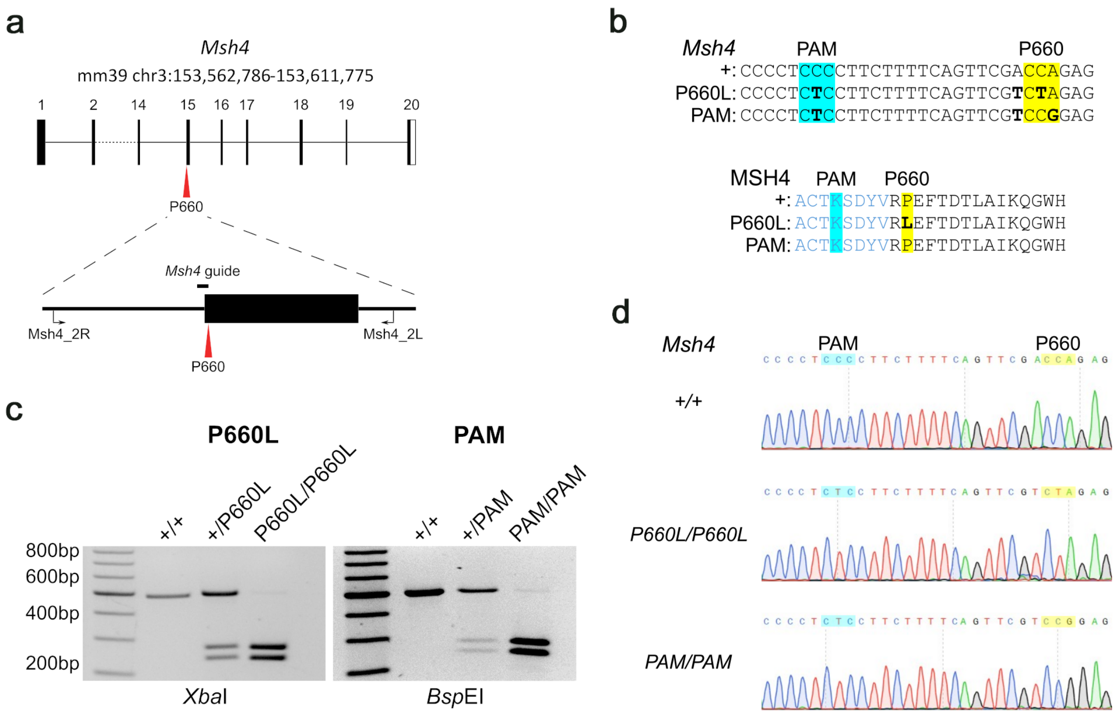


**Figure S3. (a)** Partial exon map of the mouse *Msh4* gene (transcript ENSMUST00000005630.11), showing the location of the P660 residue relative to the exon and intron borders. Exon numbers and the location of the MSH4 genotyping primers and the CRISPR/Cas9 guide are also indicated. **(b)** Nucleotide and amino acid sequences of the +, P660L and PAM *Msh4* alleles. The P660 residue is highlighted in yellow, the PAM site highlighted in blue, and nucleotide/amino acid substitutions shown in bold. Amino acids from different exons are coloured blue and black. **(c)** Example of PCR genotyping mice from the *Msh4* P660L and PAM mouse lines using primers indicated in panel c followed by restriction digest. The P660L allele includes an *Xba*I restriction site, the PAM allele includes a *Bsp*EI restriction site. Fragment sizes; no restriction site present = 484 bp, *Xba*I = 260, 224 bp, *Bsp*EI = 260, 224 bp. **(d)** Examples of DNA sequencing chromatograms of the *Msh4* genotyping PCR product from *Msh4*$^{+/+}$, *Msh4*$^{P660L/P660L}$and *Msh4*$^{PAM/PAM}$ mice. The P660 residue is highlighted in yellow, the PAM site highlighted in blue.

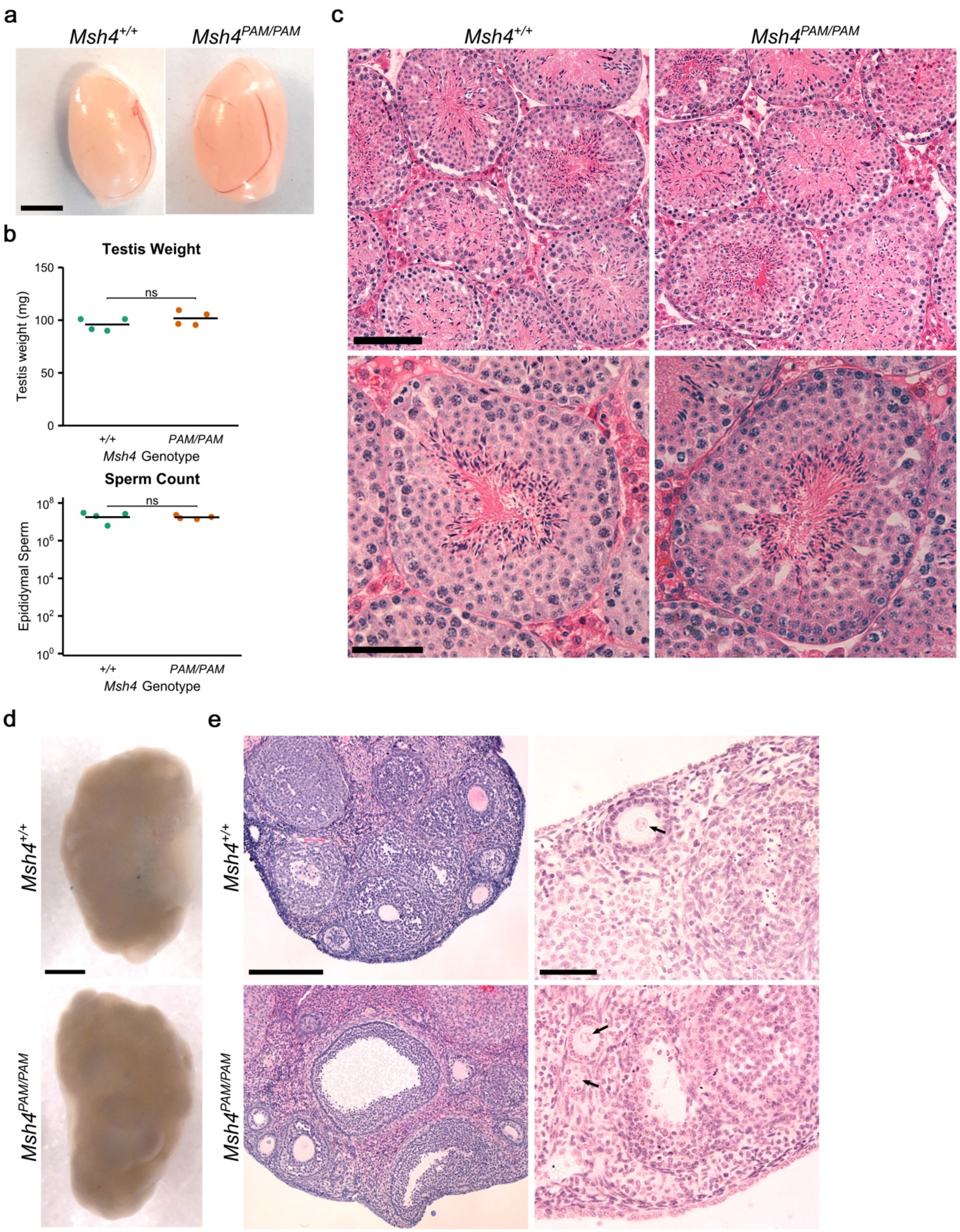


**Figure S4. (a)** Example images of *Msh4*$^{+/+}$ and *Msh4*$^{PAM/PAM}$ testes. Scale bars, 5 mm. **(b)** Testes weights and epididymal sperm counts from *Msh4*$^{+/+}$ and *Msh4*$^{PAM/PAM}$ mice. Horizontal bars indicate means. ns, not significant (Student's t-test). n = 4 mice. **(c)** Haematoxylin and eosin staining of testes sections. Scale bars, top = 100 µm and bottom = 50 µm. **(d)** Images of

*Msh4*$^{+/+}$ and *Msh4*$^{PAM/PAM}$ ovaries. Scale bar, 1 mm. **(e)** Haematoxylin and eosin staining of *Msh4* ovary sections, showing both overall tissue structure (left) and individual follicles (right). Examples of oocyte nuclei in primary or secondary follicles are indicated with arrows. Images are representative of ovaries from 4 mice per genotype. Scale bars, left = 200 µm and right = 50 µm. Table S1 provides source data for graphs in panel S4b.

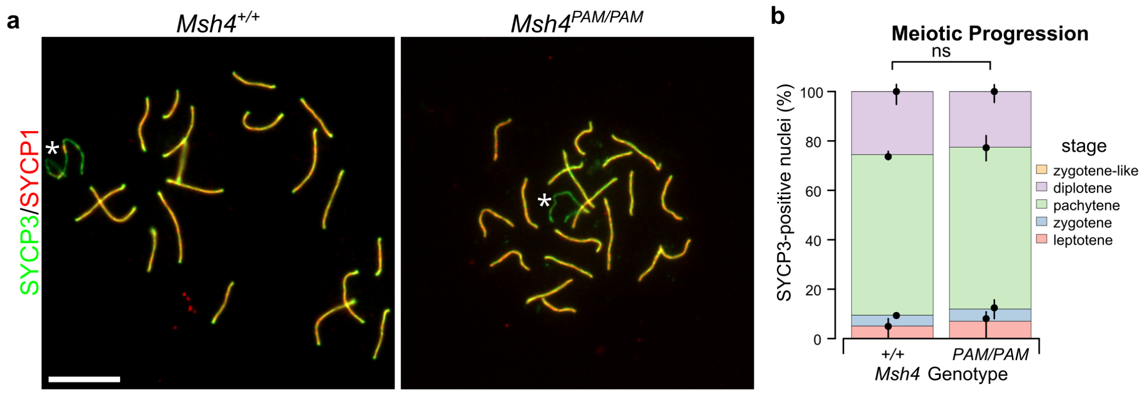


**Figure S5 (a)** Widefield imaging of pachytene *Msh4*$^{+/+}$ and *Msh4*$^{PAM/PAM}$ spermatocytes meiotic chromosome spreads immunostained for the synaptonemal complex components SYCP3 (green) and SYCP1 (red). Sex chromosomes are indicated with an asterisk. Scale bar= 10 µm. **(b)** Quantitation of percentage of total SYCP3-positive nuclei observed at each stage of prophase or in a zygotene-like state. n = 4 mice per genotype; 137 and 142 nuclei scored in total; ns, not significant (Bayesian GLMM). Table S1 provides source data for graph in panel S5b.

**Supplementary tables**

**Table S1: *C. elegans* strains used in this study**

| Strain | Genotype | Origin |
|---|---|---|
| N2 Bristol | Wild type | CGC |
| AV308 | *him-14(it21)/mnC1 [dpy-10(e128) unc-52(e444)] II* | Zalevsky et al.,1999 |
| ATG332 | *cosa-1(fq42[cosa-1::HA]) III* | Castellano-Pozo et al., 2020 |
| ATG706 | *him-14(fq179[him-14 P546L]) II/ mnC1[dpy-10(e128) unc-52(e444)] II* | This study |
| ATG800 | *him-14(fq214[him-14 G557R]) II/ mnC1[dpy-10(e128) unc-52(e444)] II; dpy-10(cn64) II* | This study |
| ATG796 | *him-14(fq211[him-14 F497C]) II* | This study |
| ATG728 | *him-14(fq179[him-14 P546L]) II /mnC1[dpy-10(e128) unc-52(e444)] II; cosa-1(fq42[cosa-1::HA]) III* | This study |

**Table S2: *C. elegans* CRISPR oligos used in this study**

| Allele | gRNA | Donor oligonucleotide for HR | Genotyping oligonucleotides |
|---|---|---|---|
| *him-14*$^{P546L}$ | GAAAATGATGGTCCAAATGT | TTGCAACTTATTCGGATCTACGTGACACATGTAAATTAACGTTCGGGCCTTCTTTTTCAATTAGTCAAGGACGTCATCCTATCCTGGA | TTCGCGCACCTAATATTCCA<br><br>AAAGAAGGCCCGAACGTTAA |
| *him-14*$^{G557R}$ | TCAAGGACGTCATCCTATCC | AAACCTACATTTGGACCATCATTTTCAATTAGTCAGCGTCGACACCCCATTTTAGACTGGGATGATTCCGAGAAAACAATCACAAACGA | ACTCGTCGGTTCGTCTTTCT<br><br>CTGTGCCATGATTGCCAGTT |
| *him-14*$^{F497C}$ | AGTTGTAGCTGAAATGTTTT | TTTATGATATTATAAATTGCAGATCGGCTAGAGCAGGTCGTTGCCGAGATGTGTTTAGCTAGCGATGTAATTGTGTGTGATATGATTGAAGAAATGCA | ACTCGTCGGTTCGTCTTTCT<br><br>CTGTGCCATGATTGCCAGTT |

**Table S3: Mouse CRISPR oligos used in this study**

| Name | Type | Sequence |
|---|---|---|
| Msh4guide | crRNA | CTGGTCGAACTGAAAAGAAG |
| Msh4_P660L | ssDNA repair oligo | AAGAATGGGATGCCATCCTTGTTTGATTGC<br>TAAAGTGTCAGTAAACTCTaGaCGAACTGA<br>AAAGAAGGaGAGGGGTTAAAGCTGAATGTA<br>TTTATATTTAAATATATTGTGGTGTGCACA |
| Msh4_PAM | ssDNA repair oligo | AAGAATGGGATGCCATCCTTGTTTGATTGC<br>TAAAGTGTCAGTAAACTCcGGaCGAACTGA<br>AAAGAAGGaGAGGGGTTAAAGCTGAATGTA<br>TTTATATTTAAATATATTGTGGTGTGCACA |
| Msh4_2L | Genotyping primer | GCCTATGCGAGTGGTTCTTG |
| Msh4_2R | Genotyping primer | CCAGGGCAGTGAAAAGGAAC |
